# Single cell proteomic analysis defines discrete neutrophil functional states in human glioblastoma

**DOI:** 10.1101/2025.07.23.666094

**Authors:** Pranvera Sadiku, Alejandro J. Brenes, Rupert L. Mayer, Leila Reyes, Patricia Coelho, Gabi van Stralen, Ailiang Zhang, Manuel A. Sanchez-Garcia, Emily R. Watts, Imran Liaquat, Andrew J.M. Howden, Ikeoluwa Adekoya, Anuka Boldbaatar, Allan MacRaild, Sarah Risbridger, Gillian M. Morrison, Heather MacPherson, Caroline Bruce, Shonna Johnston, Robert Grecian, Fiona A. Murphy, Steven M. Pollard, Paul M. Brennan, Karl Mechtler, Sarah R. Walmsley

## Abstract

Neutrophils are vital innate immune cells shown to infiltrate glioblastomas, however we currently lack the molecular understanding of their functional states within the tumour niche. Given that neutrophils are known to display a prominent discordance between mRNA and protein abundance, we developed ultra-sensitive mini-bulk and single cell proteomic (SCP) workflows to study the heterogeneity of peripheral blood and tumour associated neutrophils (TAN) from patients with glioblastoma. Mini-bulk analysis enabled a deeper protein coverage of circulating immature, mature and TAN populations, defining signatures of maturity and demonstrating that TANs resemble mature circulating neutrophils. Analysis of the SCP data resulted in the detection of >1,100 proteins from a single TAN providing a detailed characterization of neutrophil subsets in glioblastoma. Our approach shows evidence of pathogenic and anti-tumorigenic clusters and discovers cell states invisible to scRNAseq, opening new opportunities to selectively target pro-tumoural neutrophil states.

## MAIN

Glioblastoma (GBM) is a grade IV glioma, the most common and aggressive primary brain cancer in human adults^1^. The current standard of care for affected patients consists of treatment involving a combination of maximal surgical resection with radio-and chemotherapy. Despite treatment, prognosis is very poor with a 5-year patient survival rate of 4%^2^. With extensive neutrophil infiltration into GBM tumours reported, neutrophils have the potential to present themselves as potent therapeutic targets^3^. To effectively target neutrophils in GBM we firstly need to understand the relative functional contribution of these different neutrophil populations within the tumour site.

Alongside their conventional antimicrobial roles, neutrophils have emerged as important regulators of cancer. Their function remains paradoxical however, with both pro- and anti-tumourigenic effects reported at multiple stages of disease^4,5^. Neutrophils are short lived rapidly turned over cells with the bone marrow producing more than 100 billion per day in homeostasis^6^. They display significant transcriptional and functional heterogeneity in circulation, primary and metastatic tumour sites. This complexity is further amplified by cancer associated perturbations in myelopoiesis^5,7^. Single cell transcriptional analysis has been transformational in defining the presence of these different neutrophil states, and more recently has been used to predict a terminal transcriptional state in the tumour niche^8^. However, changes in rates of protein synthesis versus degradation^9^ and the long-term storage of proteins within granules together mean that transcriptomes cannot always predict effector functions. This is particularly relevant in both mature and immature neutrophil populations where weak mRNA to protein correlation is observed^10^. Our objective, therefore, was to develop a more comprehensive proteomic platform to enable us to define neutrophil subsets by function using high resolution mass spectrometry (MS).

MS-based proteomics has proven to be a highly insightful tool to define neutrophil phenotypes in humans^10–14^. However, these bulk proteomic workflows required millions of cells for the analysis to be viable, limiting their use to peripheral blood neutrophils. Recent progress on sample processing^15–19^, instrument sensitivity^20–22^ and software tools^23,24^, have facilitated the study of fewer cells^25,26^, as well as enabling the proteomic characterisation of single cells. A new generation of mass spectrometers has significantly increased sensitivity and empowered single cell proteomics (SCP) to identify >5,000 proteins per single HeLa cell^20,21^. This remarkable improvement has opened the possibility to study the functional heterogeneity of primary human cells composed of low protein content. It is important to note that neutrophils contain a much lower total protein mass, estimated to be less than 60pg per cell, compared to HeLa cells, estimated to contain 250pg of protein per cell.

Here we combine flow cytometry and MS-based proteomic workflows to study rare neutrophil populations, both from the peripheral blood and from glioblastoma (GBM) tumours. We perform a mini-bulk (500 cells) proteomic analysis of circulating mature CD10+, immature CD10-low density neutrophils (LDN) and normal density neutrophils (NDNs) as well as GBM tumour associated neutrophils (TAN), finding that TANs much more closely resemble mature CD10+ neutrophil populations, but with increased mitochondrial content. We performed ultra-sensitive SCP analysis of TANs to define functional neutrophil clusters within the GBM tumours. The power to identify > 1,100 proteins per single neutrophil provides us the ability for the first time to define the presence of discrete neutrophil functional clusters which we assign as armed, engaged, vital NETs, exhausted, lytic NETs, immunosuppressive and angiogenic and vascular immature. The added value of SCP has enabled the characterization of functional states not previously captured by single cell RNA sequencing (scRNAseq), as a result of imperfect correlation between the tissue neutrophil transcriptome and proteome. Consequently, our work stratifies neutrophil heterogeneity by effector function and opens new avenues to engage anti-tumoural neutrophil responses for GBM disease whilst also providing a novel platform for the study of neutrophils in disease at the single cell resolution.

## RESULTS

To study rare neutrophil subsets in GBM, we developed mass spectrometry-based proteomics methods to work on low cell numbers. Similar to previous work^25^, we established a mini-bulk method (500 cells) and a SCP method, both optimized for human neutrophils. Mini-bulk was used to analyze both peripheral blood neutrophils as well as GBM TANs, while only TANs were analysed with SCP. Both blood and tissue neutrophils were sorted by flow cytometry directly into 384 well plates containing the master mix (Extended Data Fig. 1A-C). Plates were stored at minus -80°C, posteriorly processed on the cellenONE X1 and analysed on an Orbitrap Astral equipped with a FAIMS Pro Duo interface using a 50 sample per day (SPD) method (Fig.1A). This workflow enabled deeper coverage from just 500 cells and the first single cell proteomic characterization of human neutrophils.

**Fig. 1.**
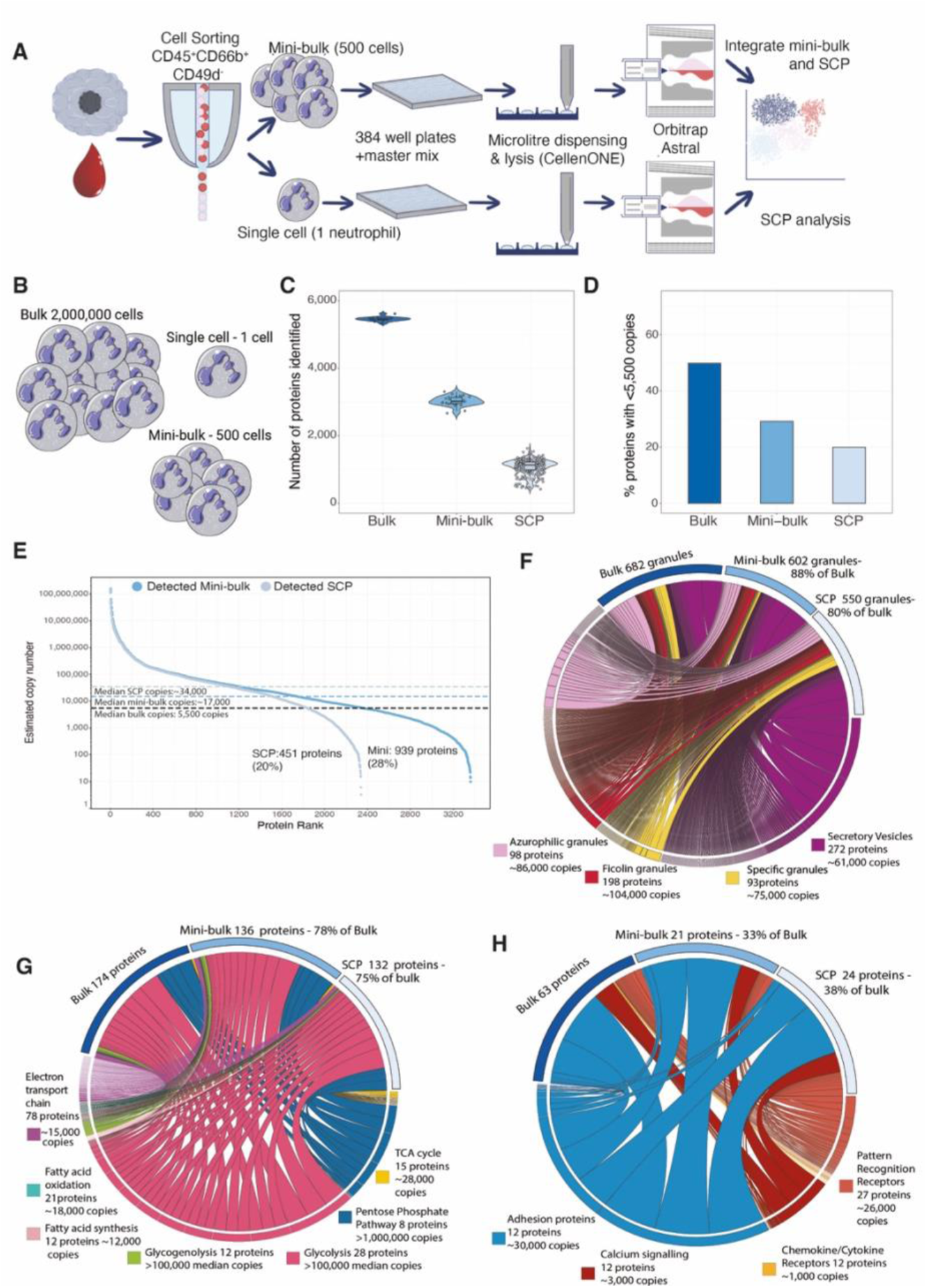
Low cell number mass spectrometry-based proteomic workflows. **(A)** Schematic showing the mini-bulk and single cell proteomic experimental design. Neutrophils (CD45+, CD66b+, CD49d-) were sorted using flow cytometry directly into 384 well plates containing the master mix, later processed on the cellenONE X1 and analysed on the Orbitrap Astral. (**B**) Schematic showing the number of cells that are analysed in bulk, mini-bulk and single cell proteomics. (**C**) Boxplot showing the number of proteins identified in bulk (n=15), mini-bulk (n=29) and single cell proteomics (n=277). **(D)** Percentage of proteins identified with less than 5,500 copies (median copy number in bulk) across all 3 proteomic workflows. Dot plot showing proteins identified in **(E)** mini-bulk and SCP ordered by their median copy number in bulk. Chord diagrams showing the coverage of **(F)** granule proteins, **(G)** metabolic proteins and **(H)** immune signaling proteins across bulk, mini-bulk and single cell proteomics. For all chord diagrams the ribbons are sized by median copy number of each protein. For all boxplots, the top and bottom hinges represent the 1st and 3rd quartiles. The top whisker extends from the hinge to the largest value no further than 1.5 × interquartile range (IQR) from the hinge; the bottom whisker extends from the hinge to the smallest value at most 1.5 × IQR of the hinge. SCP, single cell proteomics; TCA, tricarboxylic acid cycle.

We compared the protein identification of our two optimized low cell number methods to the standard bulk workflows which required 2 million neutrophils per sample (Fig.1B). The data from all 3 proteomic methods were searched using Spectronaut 19 (see methods) with a human SwissProt database that included isoforms. Details on the parameters are expanded in the methods section. As expected, the bulk proteomic workflow enabled the most comprehensive coverage of the neutrophil proteomes, with >5,700 protein groups (proteins) identified per sample. However, in the bulk workflow a total 1.5 µg of peptides were injected into the mass spectrometer. The mini-bulk analysis enabled the identification of >3,000 proteins per sample from just 500 cells, with the equivalent of ∼15ng injected into the mass spectrometer. Finally, our SCP workflow enabled the identification of >1,100 proteins per single neutrophil after tissue dissociation and cell sorting (Fig.1C) from an estimated ∼30pg of peptides injected. The mini-bulk and SCP workflows achieved remarkable coverage considering the low input.

Leveraging the bulk data, we calculated the estimated copy numbers using the proteomic ruler^27^ (see methods) and studied proteins identified across all 3 workflows. The median estimated copy number of all proteins identified in bulk was ∼5,500, thus 50% of proteins identified in bulk were present with fewer than 5,500 estimated copies (Fig.1D). Mini-bulk identified fewer low abundance proteins, with only 28% of the proteins identified in mini-bulk estimated to have <5,500 copies based on the bulk data (Fig.1D) and the median copy number of these proteins being ∼17,000 copies (Fig.1E). In SCP only 20% of proteins identified were estimated to have <5,500 estimated copies based on the bulk data (Figure 1D&E) and the median copy number being ∼34,000 copies (Fig. 1E). In summary low abundance proteins were challenging to detect with low cell number workflows, however even SCP identified 20% of the proteins detected with <5,500 copies in bulk.

We were interested to examine the proteomic coverage across all 3 methods of proteins that are vital to neutrophil function, focusing on neutrophil granules, metabolic proteins as well as proteins involved in immune signaling pathways. Our data showed that neutrophil granule proteins are highly abundant, with azurophilic granules (AG), specific granules (SG), ficolin granules (FG) and secretory vesicles (SV) all having a median copy number >60,000 copies. In bulk we could detect 682 granules, of which 88% were also detected by mini-bulk and 80% by SCP (Fig.1F), showing a remarkable capacity for SCP to characterize important neutrophil effector proteins. We next studied metabolic proteins which are vital to fuel neutrophil effector functions. The metabolic proteins were considerably less abundant than granule proteins but still displayed a median of >15,000 copies. 78% of the proteins identified by bulk were present in the mini-bulk analysis with excellent coverage (75% of bulk proteins) also retained within the SCP data (Fig. 1G). We next focused on important neutrophil effector function regulators, such as adhesion proteins, proteins required for calcium signaling responses, cytokine and chemokine receptors and pattern recognition receptors (Fig. 1H). The adhesion proteins were highly abundant and showed good coverage, but low abundance chemokine receptors were difficult to detect even in bulk, and most were not detected with mini-bulk or SCP, showing the current limitations to identify important low abundance proteins.

### Mini-bulk enables the analysis of mature and immature blood neutrophils and TANs

To define GBM neutrophil subsets by cell surface marker expression, we first undertook flow cytometry analysis of peripheral blood from healthy control (HC) and GBM patients. GBM patients had a significantly elevated leukocyte count (Extended Data Fig. 2A) and altered proportions of lymphocytes and neutrophils (Extended Data Fig. 2B). This was a consequence of a significantly higher number of neutrophils (Extended Data Fig. 2C) and a lower lymphocyte count resulting in an increased neutrophil to lymphocyte ratio (NLR) in GBM peripheral blood compared to healthy controls (Extended Data Fig. 2D). Expansion of a circulating low density neutrophil subset composed of a mixture of mature and immature cells has previously been reported in lung and breast cancer patients with distinct phenotypic properties^28^. We detected the presence of an immature CD10 negative neutrophil subset in the blood of GBM patients which was significantly increased in the LDN fraction (Extended Data Fig. 2E, F, G). Furthermore, this immature subset had features of activation with high CD66b and reduced CD11b expression (Extended Data Fig. 2H, I).

Given the idenitification of multiple GBM neutrophil subsets by our flow cytometry analysis, we decided to further characterize these populations using our mini-bulk method. We stratified the neutrophils by density and maturity using CD10, with CD10+ representing mature neutrophils and CD10-immature neutrophils. Blood low density neutrophils (LDN) CD10-, blood LDN CD10+, blood normal density neutrophils (NDN) CD10+ and CD45+CD66b+CD49d-TAN derived from 6 GBM patients and blood NDN CD10+ neutrophils from 6 healthy control donors were analysed (Figure 2A). Across all 5 neutrophil populations it was possible to identify >3,000 proteins per 500 cell sample (Fig. 2B) with a median of >14,500 peptides (Fig. 2C), showing the robustness of the method. Relative intensity based absolute quantification (riBAQ) was used to normalise and analyse the mini-bulk data (see methods). An initial principal component analysis (PCA) of the riBAQ data displayed unexpected results. The neutrophil populations displayed no clear sepparation based on density, with LDN and NDN CD10+ groups showing limited separation (Fig. 2D). TANs also clustered within the CD10+ section and it was only the CD10-negative population that showed a clear segregation. Studying the top 25 proteins that influenced the PCA revealed ribosomal proteins and PCNA marking this separation (Extended Data Fig. 3A).

**Fig. 2.**
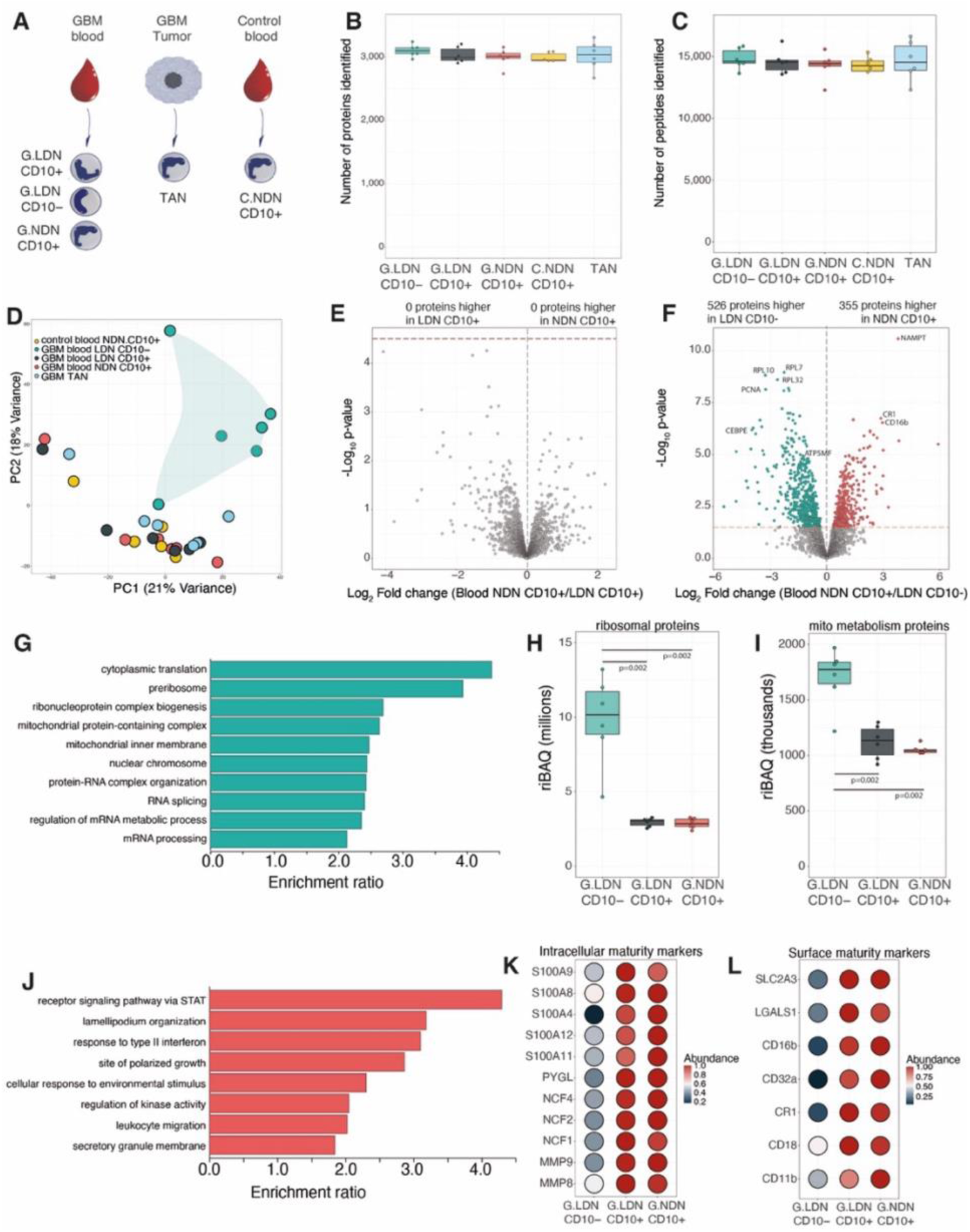
Mature (CD10+) and immature (CD10-) neutrophil proteomes. **(A)** Schematic showing the experimental design for mini-bulk proteomic analysis of human circulating blood and tumour associated neutrophils (TAN). (**B**) Number of proteins identified in the mini-bulk analysis (Control NDN CD10+ n=5, for all other conditions n=6) (**C**) Number of peptides identified in the mini-bulk analysis (Control NDN CD10+ n=5, all others n=6). **(D)** Principal component analysis (PCA) of the mini-bulk proteomic samples (n=29). Volcano plots comparing **(E)** CD10+ normal density neutrophils (NDN) to CD10+ low density neutrophils (LDN) and (**F**) CD10+ NDN to CD10-LDN. (**F**) GO enrichment analysis for proteins significantly increased in abundance in CD10-LDN. Boxplots (n=6 across all conditions) showing the summed protein abundance of **(H)** ribosomal proteins, **(I)** mitochondrial metabolism proteins across LDN CD10-, LDN CD10+, NDN CD10+. **(J)** (**F**) GO enrichment analysis for proteins significantly increased in abundance in CD10+NDN. Dot plot showing **(K)** intracellular maturity markers and **(L)** cell surface maturity markers across LDN CD10-, LDN CD10+, NDN CD10+. For all boxplots, the top and bottom hinges represent the 1st and 3rd quartiles. The top whisker extends from the hinge to the largest value no further than 1.5 × interquartile range (IQR) from the hinge; the bottom whisker extends from the hinge to the smallest value at most 1.5 × IQR of the hinge. All volcano plots show the p-values and fold changes. All p-values were calculated with limma using Empirical Bayes statistics for differential expression. All points above the red line have a q-value<0.05

We performed a direct comparison between CD10+ LDN and NDN and found there were no proteins significantly changed between the two populations (Fig. 2E), with all granule subsets showing no difference between mature LDN and NDN (Extended Data Fig. 3B-E), suggesting density is not the primary axis of proteomic variation in neutrophils. We next compared NDN CD10+ to immature LDN CD10- and found 29% of proteins being significantly different between the two populations (Fig. 2F). A total of 526 proteins were significantly higher in abundance in the LDN CD10immature neutrophils. A gene ontology (GO) analysis of these proteins revealed they were enriched in translational, DNA and mitochondrial related GO terms (Fig. 2G). The immature neutrophils displayed >350% increased abundance of ribosomal proteins compared to mature LDN and NDN (Fig. 2H), suggesting higher translational capacity. Mitochondrial metabolism proteins were also higher (>70%) in immature cells compared to mature neutrophils (Fig. 2I), as were proliferation markers such as mKI67 and PCNA and DNA replication proteins (Extended Data Fig. 3F-L).

There were 355 proteins that were significantly more abundant in mature NDNs compared to immature LDNs. A GO analysis highlighted these were enriched in processes related to cell membrane, migration and inflammatory response (Fig. 2J), suggesting the potential for superior migratory and effector capacity. A higher abundance of proteins reported to associate with increased neutrophil maturation status was observed, including core components of the NADPH oxidase, granule proteins MMP8, MMP9, S100A8, S100A9 (Fig. 2K) and the cell surface markers CD11b, CD18, CD16 and SLC2A3 (Fig. 2L).

### TANs resemble mature circulating neutrophil populations

We were interested to understand whether tumour associated neutrophil populations had a unique proteomic signature. TANs displayed higher coefficient of variation (CV) when compared to the blood derived populations (Fig. 3A). In line with the PCA overview, which displayed no clear separation of TANs from CD10+ populations, a direct comparison of TANs to mature CD10+ NDN reflected only 8 proteins significantly changed in abundance (Fig. 3B), suggesting limited differences compared to normal density mature neutrophils. Of the few proteins changed TANs displayed increased abundance of brain specific markers such as MBP and MAOB while L-selectin, the cell adhesion molecule important for trans-endothelial migration was higher in the NDN CD10+. We then performed a direct comparison of TANs to LDN CD10-, and found this revealed 553 proteins changed in abundance (Fig. 3C). Of these 553 proteins, proteins with higher abundance in TANs were enriched for GO terms relating to migration and adhesion (Fig. 3D) and demonstrated increased abundance of maturity markers (Extended Data Fig. 4A-H). Proteins decreased in abundance in TANs related to ribosomal and DNA replication proteins (Fig. 3E-G). Markers of proliferation were also significantly lower in TANs compared to the immature cells (Extended Data Fig. 4I&J). Together these highlight that TANs are much more similar to CD10+ neutrophil populations than the immature CD10-neutrophils. Interestingly, TANs did however, display an intermediate mitochondrial phenotype, when compared to CD10-LDN and CD10+ NDN (Fig. 2H&I), suggesting the potential for mitochondrial metabolic adaptation within the tumour niche.

**Fig. 3.**
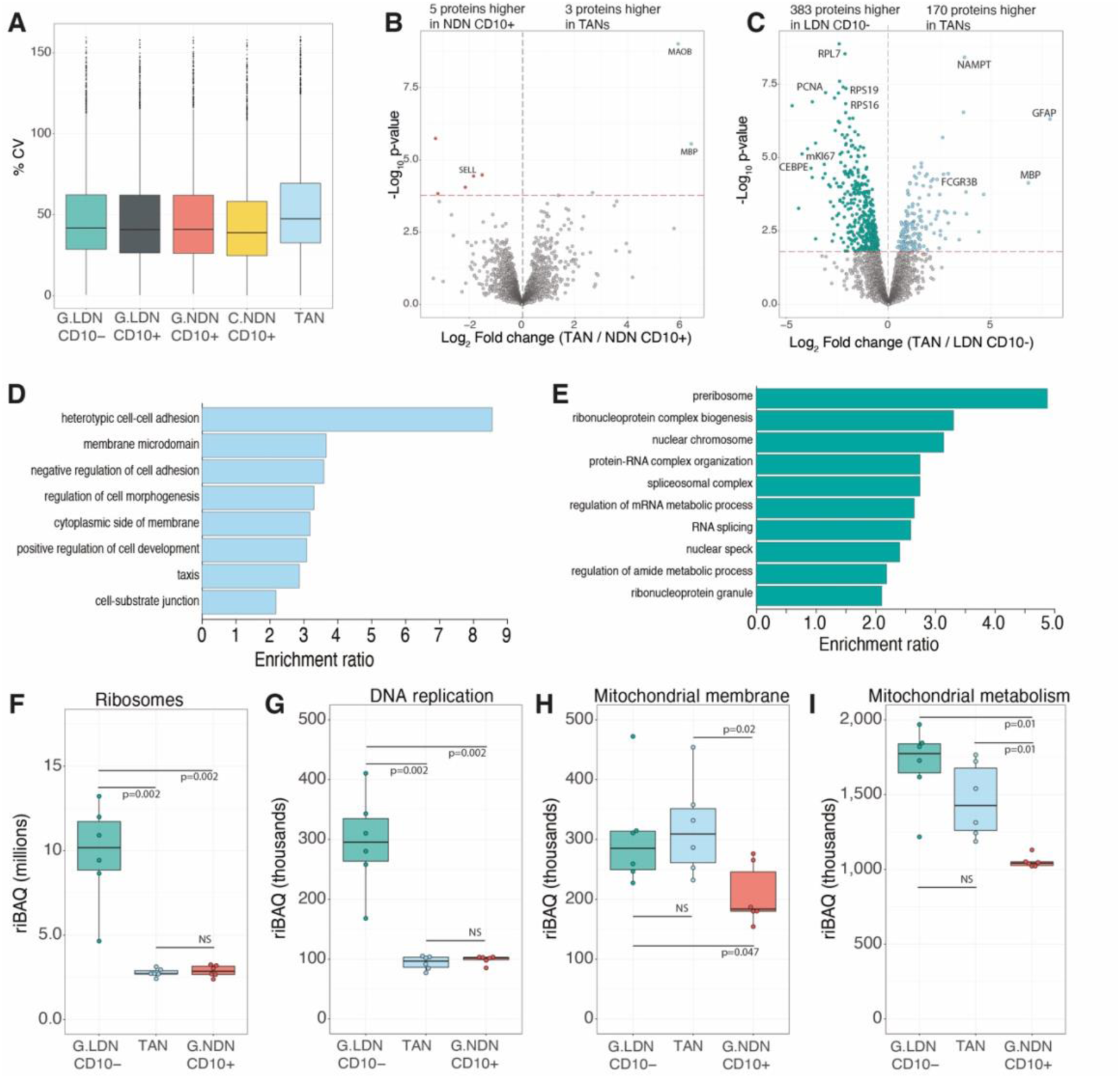
TANs resemble mature neutrophils with a mitochondrial phenotype. **(A)** Boxplot showing the coefficient of variation (CV) for all mini-bulk populations. Volcano plots comparing **(B)** CD10+ normal density neutrophils (NDN) to tumour associated neutrophils (TANs) and (**C**) CD10-low density neutrophils (LDN) to TANs. GO enrichment analysis for proteins **(D)** significantly increased in abundance in TANs compared to LDN CD10- and **(E)** significantly decreased in abundance in TANs compared to LDN CD10-. Boxplots (n=6 across all conditions) showing the summed riBAQ for all **(F)** ribosomal proteins, **(G)** DNA replication proteins, **(H)** mitochondrial membrane proteins and **(I)** mitochondrial metabolism proteins. For all boxplots, the top and bottom hinges represent the 1st and 3rd quartiles. The top whisker extends from the hinge to the largest value no further than 1.5 × interquartile range (IQR) from the hinge; the bottom whisker extends from the hinge to the smallest value at most 1.5 × IQR of the hinge. All volcano plots show the p-values and fold changes. All p-values were calculated with limma using Empirical Bayes statistics for differential expression. All points above the red line have a q-value<0.05. All protein family p-values were calculated using Welch’s t test.

### Single cell proteomics identifies distinct neutrophil states in the Glioblastoma

To investigate neutrophil heterogeneity in GBM, we utilized ultra-sensitive MS-based single cell proteomics (SCP). We optimized a workflow where GBM TAN were FACS sorted into 384 well plates and analysed them on the Orbitrap Astral with FAIMS Pro using a 50 sample per day workflow (see methods). For this analysis the mini-bulk proteomic data was integrated to deconvolute the SCP data sets for specific signatures (Figure 1A) and immature populations within the SCP data were identified using the maturity markers that were identified in the NDN CD10+ to LDN CD10-comparison.

A total of 330 neutrophils derived from tumours of the same 6 patients used in the blood and tumour mini-bulk analysis were processed. As previously mentioned, the MS-based raw files were searched with Spectronaut v19.4, using software parameters with increased stringency compared to the default settings (see methods). From the results, cells with less than 400 proteins identified were filtered out (see methods), leaving a total of 277 cells for further analysis (Extended Data Fig. 5A-F). The intensity data produced by Spectronaut was normalized using a riBAQ like approach (see methods) and the data analysed in Seurat with batches integrated using harmony (see methods). Across the 277 cells a median of > 1,100 proteins per single neutrophil were identified (Figure 1B). The preliminary analysis of the riBAQ like intensity data across the 2,000 most variable protein features (see methods) in Seurat led to the identification of seven distinct neutrophil populations (Fig. 4A), which we assign as (1) armed, (2) engaged, (3) vital NETing, (4) exhausted, (5) immunosuppressive and angiogenic, (6) lytic NETing and (7) vascular immature clusters. The most abundant was the armed neutrophil subset, and the least abundant the vascular immature subset (Fig. 4B).

**Fig. 4.**
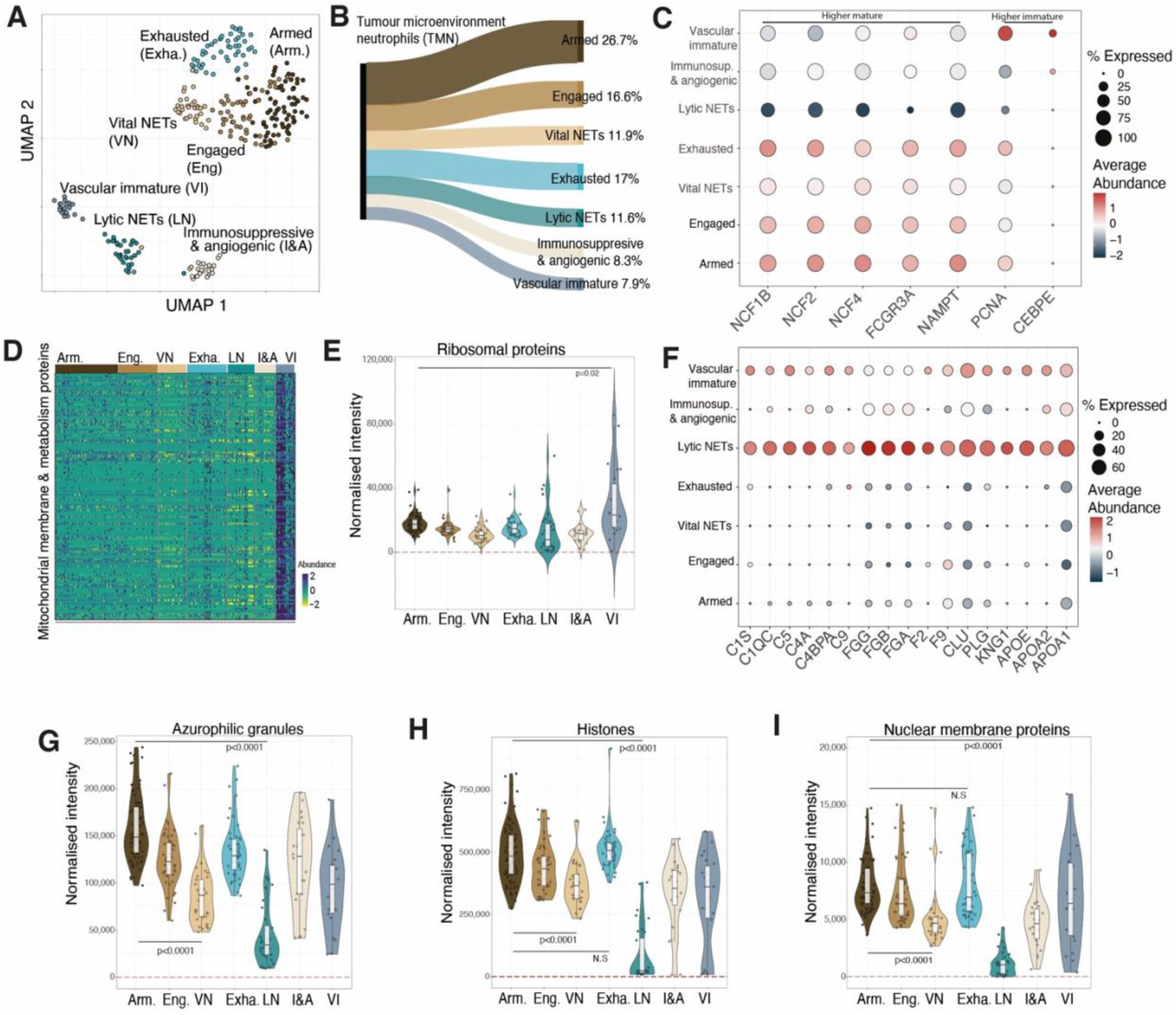
Single cell proteomic analysis of TANs identified discrete functional clusters. **(A)** UMAP showing the TAN clusters. (n=277) **(B)** Sankey diagram showing the proportions represented by each cluster. **(C)** Dotplot showing maturity marker proteins across all clusters. **(D)** Heatmap showing mitochondrial metabolism and mitochondrial membrane proteins across all cells ordered by cluster. **(E)** Boxplot showing the summed normalised intensity of all ribosomal proteins. **(F)** Dotplot showing complement and coagulation proteins across all clusters. Boxplots showing the summed normalised intensity of all **(G)** histones, **(H)** nuclear membrane proteins and **(I)** azurophilic granules across all clusters. Across all boxplots Armed (n=76), Engaged (n=46), Vital NETs (n=33), Exhausted (n=47), Lytic NETs (n=32), Immunosuppressive and angiogenic (n=23) and Vascular Immature (n=22). For all boxplots the top and bottom hinges represent the 1st and 3rd quartiles. The top whisker extends from the hinge to the largest value no further than 1.5 × interquartile range (IQR) from the hinge; the bottom whisker extends from the hinge to the smallest value at most 1.5 × IQR of the hinge. Protein family boxplot p-values were calculated using Welchs T-test.

### Vascular signature of lytic NETosis and immature GBM populations with reduced frequency of immature neutrophils in the tumour niche

To assess the contribution of immature neutrophils to the tumour neutrophil pool we integrated the protein level maturity markers identified in the mini-bulk data with the SCP and identified one population, representing 8% of the cells, displaying an immature signature (Fig. 4C). This population, labelled as vascular immature, was defined by increased abundance of PCNA and CEBPE (Fig. 4C) with concurrent increased abundance of mitochondrial metabolic enzymes including PDHB, IDH3A and ACAT1, mitochondrial membrane proteins including TOMM5 and IMMT (Fig. 4D), and overall ribosomal proteins including RPL4, RPL7 and RPL15 (Fig. 4E). The combined signature closely resembled the LDN CD10-population identified by our mini-bulk analysis. This immature tumour neutrophil population was reduced in frequency compared to the CD10^-^ peripheral blood compartment (37 %) identified by flow cytometry (Figure 2). Within this immature tumour population we also identified a signature that suggested it was located within the vasculature, with elevated levels of complement and coagulation proteins (Fig. 4F).

A second neutrophil population was also observed to display high abundance of complement proteins, fibrinogens, F2 and F9 coagulation factors, clusterin and plasminogen (Fig. 4F). This population displayed loss of highly abundant proteins that are extravasated during lytic NETosis. It displayed an 83% reduction in total histone proteins (Fig. 4G), an 85% reduction in nuclear membrane proteins (Fig. 4H) and an 80% reduction in the abundance of primary azurophilic granules (Fig. 4I). Similar pronounced reductions were observed in secondary granules, ficolin granules and secretory vesicles (Extended Data Fig. 6A-C). In keeping with a cell undergoing lytic NETosis these neutrophils had also lost their protein capacity to perform effector functions or respond to extra-cellular signals with a sharp reduction in metabolic enzymes required to fuel neutrophil effector functions (Extended Data Fig. 6D) and cell membrane proteins required to respond to DAMPs and PAMPS (Extended Data Fig. 6E).

### Neutrophils degranulate and undergo vital NETosis within the GBM tumour

Of the mature neutrophil populations present within the tumour, SCP identified cell states that correlated with a functional trajectory. The most abundant population, the armed neutrophils, have a signature consistent with recent extravasation with the highest abundance of cytoskeletal and motility related proteins (Fig. 5A) including ITGB2 (CD18) and ITGAM (Cd11b), vital integrins forming the MAC1 complex. They also display high histone abundance (Fig. 5B) and the highest abundance of granule proteins (Fig. 5C). Engaged neutrophils demonstrate signs of granule release, exemplified by reductions in myeloperoxidase (MPO) and calprotectin subunit S100A8 content, and a trajectory towards vital NETosis (Fig. 5 D&E) where loss of granule proteins and secretory vesicles (Fig. 5F) is associated with a reduction in histones (Fig. 5G&H) and nuclear and cell membrane proteins (Fig. 5I).

**Fig. 5.**
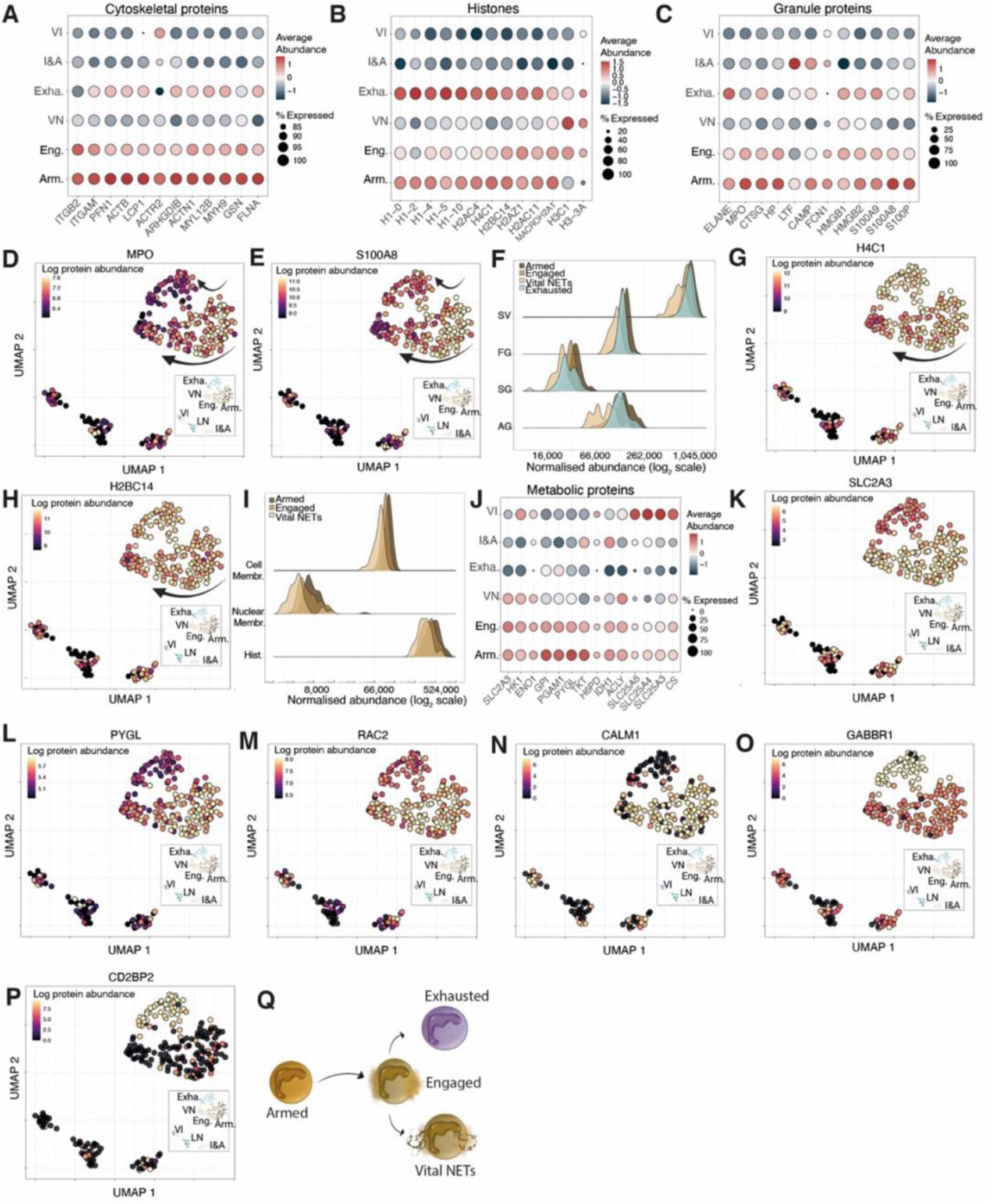
Predicted trajectory between neutrophil functional states. Dotplots for **(A)** cytoskeletal proteins, **(B)** histones and **(C)** granule proteins across armed (Arm.), engaged (Eng.), vital NETs (VN), exhausted (Exh), immunosuppressive and angiogenic (I&A) and vascular immature neutrophils (VI). UMAPs for **(D)** myeloperoxidase (MPO) and **(E)** S100 calcium-binding protein A8 (S100A8). **(F)** Ridgeplot for azurophilic granules (AG), specific granules (SG), ficolin granules (FG) and secretory vesicles (SV) across armed, engaged, vital NETs and exhausted clusters. UMAPs for **(G)** H4 Clustered Histone 1 (H4C1) and **(H)** H2B Clustered Histone 14 (H2BC14). **(I)** Ridgeplot of histones, nuclear membrane proteins and cell membrane proteins across armed, engaged and vital NETs clusters. **(J)** Dotplot showing metabolism related proteins. UMAPs for **(K)** Solute Carrier Family 2 Member 3, GLUT3 (SLC2A3), **(L)** Glycogen Phosphorylase Liver form (PYGL), (**M)** Rac Family Small GTPase 2 (RAC2), **(N)** Calmodulin 1 (CALM1), **(O)** Gamma-Aminobutyric Acid Type B Receptor Subunit 1 (GABBR1) and **(P)** CD2 Cytoplasmic Tail Binding Protein 2 (CD2BP2). (Q) Schematic showing the proposed neutrophil trajectory. All Ridgeplots show the summed normalised abundance of specific protein families. For all UMAPs (n=277) the colour scale shows the log normalised protein abundance.

A discrete mature neutrophil state is also observed in which reduced expression of granule proteins and secretory vesicles is associated with retained levels of histone proteins and metabolic anergy. We have termed these exhausted neutrophils. These cells have a significant reduction in proteins involved in vital metabolic processes including glycolysis, glycogenolysis and the pentose phosphate pathway (Fig. 5J), exemplified by low level expression of the key regulatory proteins glucose transporter GLUT 3 (SLC2A3) and glycogen phosphorylase (PYGL) (Fig. 5K&L). In keeping with a reduced capacity to respond, this dysfunction also extends to regulators of neutrophil function including the NADPH oxidase regulator RAC2 (Fig. 5M), and the calcium binding protein calmodulin (Fig. 5N). Interestingly this population was also defined by high abundance of GABBR1,a Gabba receptor associated with innate lymphoid cell inhibition^29^ (Fig. 5O) and CD2BP2, a CD2 binding protein with unknown neutrophil effector functions (Fig. 5P). Together, this data suggests that neutrophils arrive armed and poised to respond, they get engaged with potential end points of exhaustion or vital NETosis (Fig. 5Q).

### Immunosuppresive & angiogenic neutrophils have a phagocytic signature

The proteomic data also uncovered the presence of a distinct neutrophil state within the tumour that displayed an immunosuppressive signature^30^, with high abundance of Arginase 1 (ARG1) and S100A7 (Fig. 6A&B). This cluster was further defined by an increase in abundance of the pro-angiogenic factors ECM1 (extracellular matrix protein 1) (Fig. 6C) and S100A10 (Fig. 6D) thus we have termed these immunosuppressive and angiogenic (I&A) neutrophils. A reduction in the matrix metalloproteinases, MMP8 (Fig. 6E) & MMP9 (Fig. 6F) was also revealed. This is of interest as neutrophil release of MMPs has also been shown to promote angiogenesis through the degradation of the extracellular matrix. In addition, immunosuppressive and angiogenic neutrophils displayed evidence of enhanced lysosomal capacity with increased abundance of core lysosomal proteins (Fig. 6G), including Cathepsin D (CTSD), lysosomal associated membrane protein 1 (LAMP1) and scavenger receptor class B member 2 (SCARB2). This was associated with a global increase in protein degradation machinery, including the core components of ubiquitin-proteasome complex (Fig. 6H), with evidence of enhanced activity provided by higher levels of free ubiquitin (Fig. 6I). In keeping with enhanced phagocytic activity this cluster also demonstrated an increase in abundance of extracellular immunoglobulin (Ig) proteins (Fig. 6J) and non-neutrophil specific proteins normally related to the desmosomes (Fig. 6K). Within the tumour niche we have therefore identified a subset of pro-tumoural neutrophils that can phagocytose and process extracellular proteins via the lysosomal compartment, activate T cell suppression pathways and promote angiogenesis.

**Fig. 6.**
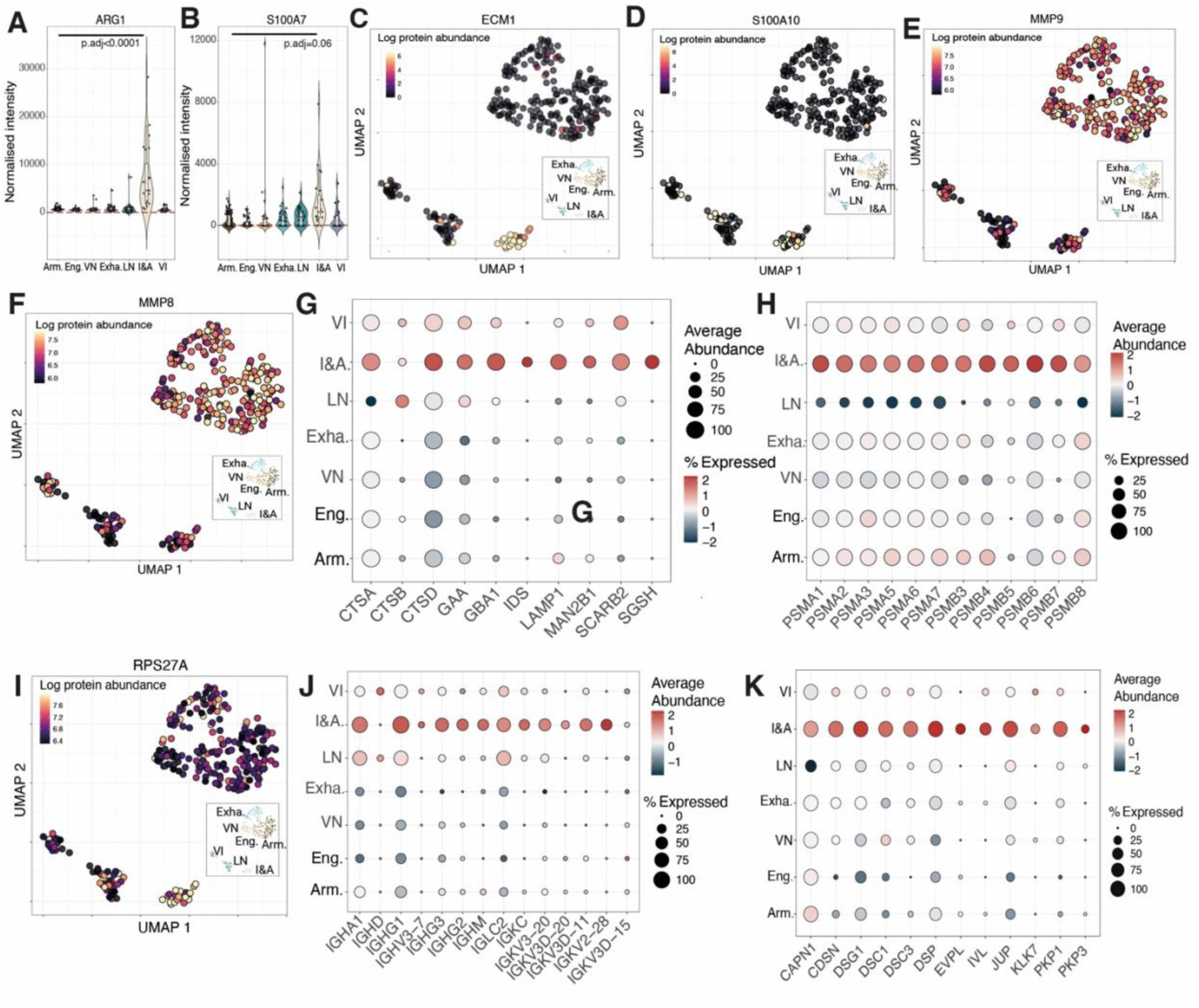
Presence of an immunosuppressive and angiogenic neutrophil subset. Boxplots for **(A)** Arginase 1 (ARG1) and **(B)** S100 Calcium Binding Protein A7 (S100A7) showing the normalised protein abundance across all clusters. UMAPs for **(C)** Extracellular Matrix Protein 1 (ECM1), **(D)** S100 Calcium Binding Protein A10 (S100A10), **(E)** Matrix Metallopeptidase 8 (MMP9) and **(F)** Matrix Metallopeptidase 8 (MMP8). Dotplots showing **(G)** lysosomal proteins and the **(H)** 20s proteasome proteins. **(I)** UMAP for the ubiquitin protein (RPS27A). Dotplot showing **(J)** immunoglobulin proteins and **(K)** Desmosome related proteins. For all UMAPs (n=277) the colour scale shows the log normalised protein abundance. Across all boxplots Armed (n=76), Engaged (n=46), Vital NETs (n=33), Exhausted (n=47), Lytic NETs (n=32), Immunosuppressive and angiogenic (n=23) and Vascular Immature (n=22). The top and bottom hinges represent the 1st and 3rd quartiles. The top whisker extends from the hinge to the largest value no further than 1.5 × interquartile range (IQR) from the hinge; the bottom whisker extends from the hinge to the smallest value at most 1.5 × IQR of the hinge. Individual protein boxplot adjusted p-values were calculated using Seurat with logistic regression.

## Discussion

Single cell RNAseq has been instrumental in helping to define neutrophil transcriptional subsets, however due to the strong discordance between mRNA and protein abundance^10^, an understanding of neutrophil functional heterogeneity in health and disease is lacking. Although the transcriptome can be used to predict cellular state, it is the proteome that is the key driver of biological function especially in post-mitotic cells such as neutrophils. The bulk proteomes of immune cells including neutrophils have over the years been extensively studied mapping cellular protein signatures and function to understand disease mechanisms and identify biomarkers. However, bulk proteomics obscures the single cell resolution necessary to capture complex biological tissue cell-state heterogeneity. Important advances in single cell proteomics including the original isobaric labelling methods^31,32^, nanodroplet sample processing techniques^33,34^, label free instrument optimisations^35,36^ and even spatial solutions^37,38^ have enabled the quantification of thousands of proteins from a single cell. Despite these developments, the lack of amplification strategies for proteins has mostly limited SCP analysis to larger cells containing a higher protein content such as blastomeres, oocytes and HeLa cells^39^. Whilst a single HeLa cell is estimated to contain between 150-250 pg of protein^40^, a single human neutrophil is estimated to contain between 30-60 pg of protein, posing a substantial challenge for SCP analysis ^13,39,40^.

To facilitate profiling of neutrophil proteomes with low protein input, we have developed a proteomics workflow optimised for low cell numbers, capable of measuring from 500 neutrophils down to a single FACS sorted human neutrophil dissociated from glioblastoma tumours. This high sensitivity workflow has enabled the detection of >3,000 proteins from 500 neutrophils and >1,100 proteins per single human TAN, with good coverage of the highly abundant neutrophil effector proteins exemplified by the granule proteins (80% SCP coverage), metabolic proteins (75% SCP coverage), and immune signalling proteins (38% SCP coverage). This workflow has allowed us to generate the first human single cell tissue neutrophil proteomic resource.

Neutrophils are highly dynamic cells with the capacity to adapt to the tissue niche. There is compelling evidence to show that in addition to re-programming within the tumour micro-environment, newly formed neutrophils and their progenitors can also be perturbed in the bone marrow long before they reach the tumour site^5,41^. We find that GBM alters circulating neutrophil with an expansion of a low density immature neutrophil population with a distinct proteomic signature. Our mini-bulk proteomic survey of 500 blood CD10-LDN, CD10+ LDN, CD10+ NDN and CD45+CD66b+CD49d-TAN using ultra-sensitive mass spectrometry revealed proteomic similarities between CD10+ blood and tumour associated neutrophils, however it also highlighted clear differences between the CD10-LDN and TAN populations. This is of interest given the significant contribution of LDN to the circulating neutrophil pool and work detailing deterministic reprogramming of both immature and mature neutrophils by the tumour niche^8^. Single cell proteomic survey of neutrophils harvested from resected glioblastoma tumours at time of surgery subsequently identified the presence of an immature neutrophil cluster based on the abundance of specific maturity markers defined by mini-bulk as well as mitochondrial and ribosomal protein content. However, this accounted for only 8% of the tumour neutrophil populations and was associated with an intra-vascular signature of complement and coagulation proteins. We therefore propose that in human GBM CD10-immature neutrophils are retained within the vasculature. It remains an open question as to whether there is either active exclusion of immature neutrophils with anti-tumoural capacity by prevention of their vascular egress or whether there is rapid transition of these cells into a terminally differentiated pro-tumoural state.

Importantly, our study also reveals the concurrent existence of neutrophil states with the potential capacity to engage either pro- (immunosuppressive, lytic NETotic) or anti- (armed, engaged) tumoural responses. Our work also highlights the value and contribution of SCP, as multiple neutrophil states uncovered by this data would be invisible to RNAseq. Specifically, NETotic neutrophils were characterized by prominent reductions in the abundance of histones and granule proteins whilst degranulating neutrophils were characterized by a reduction in granule protein content with conserved histone abundance. As granule proteins are preserved within mature neutrophils long after their corresponding transcripts are no longer present^10^, these functional clusters are only exposed when proteomic analysis is performed. SCP also enables characterization of less frequent populations, for example we were able to identify a cluster of neutrophils undergoing lytic NETosis not visible within the mini-bulk data. This is of interest given the recent description via 3D imaging in murine cancer models of tumour-elicited vascular-restricted neutrophils that promote tissue necrosis via NETosis and can be blocked to reduce metastatic spread^42^. The proteomic signature of GBM neutrophils undergoing lytic NETosis complements this description of compartment restricted neutrophil effector functions raising the important potential for targeted therapeutic intervention to limit the tissue necrosis associated with poor survival outcomes in GBM^43^.

Our study provides a platform to explore human neutrophil functional heterogeneity with single cell resolution and should be considered a starting point for the investigation of neutrophil contribution to homeostasis and disease pathogenesis. Future work will be required to understand what determines the different neutrophil states in cancers such as GBM and whether they are a consequence of exposure to local cues within the tumour niche e.g. hypoxia, specific cell-cell interactions, or a consequence of pre-determined trajectories that are manifest upon exposure to the tumour micro-environment. Further developments in the field of single cell spatial proteomics will allow us to start to address these important questions. For example, can the peri-vascular niche play a critical role in compartmentalization of these different effector responses as previously reported for GBM macrophage interactions^44^. Does neutrophil phagocytosis and clearance of cell debris and tumour antigens in the necrotic core itself engage immunosuppressive pro-angiogenic responses? How do neutrophil populations evolve during tumour progression, treatment and recurrence.

In summary, single cell proteomics now unlocks the door to defining functional changes in neutrophil subsets that are subverted in disease and amenable to therapeutic targeting. Moreover, the SCP and mini-bulk workflows presented here have the potential to be widely applicable to human leukocytes, in any tissue as well as disease setting further enhancing scientific development in the field of immune cell biology.

## METHODS

### Human participants

The collection of peripheral blood from healthy male and female control participants was approved by the Centre for Inflammation Research Blood Resource Management Committee (AMREC #15-HV-013) and recruited from the University of Edinburgh Centre for Inflammation Research Blood Resource. Exclusion criteria for healthy controls included the following: infection with any blood borne diseases, previous or current intravenous drug abuse, anaemia, blood clotting disorders, anticoagulant drug therapy, regular use of steroids and/or under the age of 16 years old. The collection of peripheral blood and tumour tissue from GBM male or female patients was approved by the Lothian NRS Bioresource, (East of Scotland Research Ethics Committee REC 1 #13/ES/0126) and recruited between January 2015 to 2025 from NHS Lothian hospitals, Edinburgh, Scotland, UK. Informed consent was obtained from all participants prior to sample collection.

### Isolation of leukocytes

Blood samples were collected in S-Monovete^®^ sodium citrate tubes (Sarstedt) or drawn by syringe and transferred into a 50 mL polystyrene Falcon™ tube containing 3.8% sodium citrate. Plasma was separated by centrifuging blood in a swinging bucket rotor at 350 × *g* for 20 min at 20°C (acceleration 1, brake 5). The platelet rich plasma upper layer was removed and dextran sedimentation was performed to separate leukocytes from erythrocytes by incubating the leukocyte/erythrocyte lower layer with pre-warmed (37°C) 6% v/v dextran at 37°C for maximum 30 min. The leukocyte rich upper layer was collected with a sterile pastette, centrifuged at 350 × *g* for 6 min at 20°C (acceleration 9, brake 9) and washed in fluorescence-activated cell sorting (FACS) buffer (2% v/v HI-FBS in Dulbecco’s phosphate buffered saline (DPBS) without Ca^2+^ or Mg^2+^) ready for phenotyping by flow cytometry.

### Isolation of human blood neutrophils

For the isolation of NDN from healthy control whole blood and NDN and LDN from whole blood of GBM patients, immunomagnetic isolation of neutrophils directly from whole blood using the EasySep™ Direct Human Neutrophil Isolation Kit (StemCell Technologies) was performed as per the manufacturer’s instructions, followed by a discontinuous Percoll™ gradient as previously described^48^. Briefly, the immunomagnetically isolated neutrophil pellet was resuspended in 49% Percoll™, carefully layered above 61% Percoll™ already layered onto 73% Percoll™ and centrifuging at 720 × *g* for 20 min at 20**°**C, with brake off and acceleration set at 1. NDN were harvested from the 61%/73% interface and LDN were harvested from the 49%/61% interface followed by washing with DPBS without Ca^2+^ or Mg^2+^ (Gibco) prior to phenotyping or sorting.

### Processing of GBM tumour tissue

Fresh human tumour tissue collected during surgery was dissected using a scalpel and exposed to enzymatic digestion by incubation in complete RPMI medium supplemented with 10% v/v HI-FBS (PAN-Biotech), 1% v/v penicillin-streptomycin (1000 U/L, Gibco), DNase (1000 U/L, Roche Diagnostics) and Collagenase Type IV (0.1% w/v, ThermoFisher Scientific) in sterile conditions at 37 ^○^C in a humified atmosphere containing 5% CO_2_ for 30 min. The tissue digest was mixed intermittently every 10 min during the 30 min incubation. The tissue digest was then passed through a 100 μm Fisherbrand™ cell strainer into a 50 mL polystyrene Falcon™ tube and this process was repeated using 70 μm and then 40 μm Fisherbrand™ cell strainers. The volume of cell suspension was adjusted to 50 mL with 0.9% saline and centrifuged at 350 × *g* for 6 min at 20^○^C (acceleration 9, deceleration 5) for cell pelleting. Hypotonic red cell lysis was then performed by first re-suspending the cell pellet in 10 mL of 0.2% w/v NaCl in sterile water followed by the addition of 10 mL 1.6% w/v NaCl and 0.1% w/v D-(+)-glucose in sterile water and centrifuged at 350 × *g* for 6 min at 20^○^C (acceleration 9, deceleration 5). The cell pellet was re-suspended in 5 mL 0.9% saline and used for flow cytometry phenotyping or fluorescence-activated cell sorting (FACS) of tumour neutrophils.

### Fluorescence-activated cell sorting (FACS) of human GBM tumour neutrophils

Human GBM tumour tissue was dissected and digested to a single cell suspension as described above. For samples required for SCP and mini-bulk proteomics, cells were first incubated with 2 ml of 1:200 diluted FC Block for 10 min at 4**°**C, followed by staining in 2 ml of an antibody cocktail containing BV421-anti-CD45 (1:200 final dilution), APC-anti-CD66b (1:57 final dilution), PE-anti-CD49d (1:200 final dilution) and PE-Cy7-anti-CD10 (1:57 final dilution) for 30 min at 4**°**C. Appropriate unstained and fluorescence minus one (FMO) controls were also generated. Following three washes with PBS w/o Ca^2+^ and Mg^2+^, 500 cells of healthy control circulating NDN CD10+ and circulating GBM normal density (NDN) CD10+, low density (LDN) CD10+ mature, immature LDN CD10-, tumour associated neutrophils (TANs) were sorted for mini-bulk proteomic analysis. 66 single GBM TANs from each patient were sorted single cell proteomic (SCP) analyses. A FACS Aria Fusion sorter *(Becton Dickinson)* was used to collect cells in 384 well plates containing 1mL of a cell lysis master mix per well (See mini-bulk and single cell proteomics sample preparation).

### Human GBM leukocyte phenotyping by flow cytometry

Healthy donor and GBM leukocytes (1 × 10^6^/test) were stained with an antibody cocktail mix in FACS buffer for 25 min at 4 ^○^C with appropriate unstained and fluorescence minus one (FMO) controls. Cells were washed and re-suspended in staining buffer and acquired using BD LSRFortessa™ flow cytometer (Beckton Dickinson), with compensation performed using BD FACSDiva™ software version 8.0 and data analysed in FlowJo version 10.2.

### Human GBM neutrophil phenotyping by flow cytometry

GBM NDN and LDN (0.1 × 10^6^/test) isolated from Percoll gradients and digested human GBM tumour tissue were stained with an antibody cocktail mix containing FITC-anti-CD66b (1:40), PE-Cy7-anti-CD11b (1:40), PerCP-Cy5.5-anti-CD49d (1:40), APC-Cy7-anti-CD10 (1:40) for 30 min at 4 ^○^C with appropriate unstained and FMO controls. Cells were washed and re-suspended in FACS buffer and acquired using BD LSRFortessa™ flow cytometer (Beckton Dickinson), with compensation performed using BD FACSDiva™ software version 8.0 and data analysed in FlowJo version 10.2.

### Bulk proteomics sample preparation

2 million cell pellets were lysed in 100 µl lysis buffer (5% SDS, 10 mM TCEP, 50 mM TEAB) and were then shaken at room temperature for 5 min at 1000 rpm, followed by boiling at 95**°**C for 5 min at 500 rpm. Samples were then shaken again at RT for 5 min at 1000 rpm before being sonicated for 15 cycles of 30 s on/ 30 s off with a BioRuptor (Diagenode). Benzonase was added to each sample and incubated at 37 °C for 15 min to digest DNA. Samples were then alkylated with the addition of iodoacetamide to a final concentration of 20 mM and incubated for 1 h in the dark at 22**°**C. Protein concentration was determined using EZQ protein quantitation kit (Invitrogen) as per manufacturer instructions. Protein digestion was performed using S-TRAP micro columns (Protifi). Proteins were digested with trypsin at 1:10 ratio (enzyme:protein) for 2 h at 47**°**C. Digested peptides were eluted from S-TRAP columns using 50 mM ed peptides were dried overnight before being resuspended in 40 µl 1% formic acid ready for analysis by data independent acquisition (DIA) mass spectrometry.

### Bulk proteomics mass spectrometry

The bulk proteomics data was acquired in DIA mode on an Orbitrap Exploris 480 (Thermo Scientific) coupled with an UltiMate 3000 RSLC nano (Thermo Scientific™). 1.5 µg of peptide per sample were injected. Two buffers were used: buffer A (0.1% formic acid in Milli-Q water (v/v)) and buffer B (80% acetonitrile and 0.1% formic acid in Milli-Q water (v/v). Samples were loaded at 10 μl/min onto a trap column (100 μm × 2 cm, PepMap nanoViper C18 column, 5 μm, 100 Å, Thermo Scientific™) equilibrated in 0.1% trifluoroacetic acid (TFA). The trap column was washed for 3 min at the same flow rate with 0.1% TFA then switched in-line with a Thermo Scientific™, resolving C18 column (75 μm × 50 cm, PepMap RSLC C18 column, 2 μm, 100 Å). Peptides were eluted from the column at a constant flow rate of 300 nl/min with a linear gradient from 3% buffer B to 6% buffer B in 5 min, then from 6% buffer B to 35% buffer B in 115 min, and finally to 80% buffer B within 7 min. The column was then washed with 80% buffer B for 4 min and re-equilibrated in 3% buffer B for 15 min. Two blanks (1% formic acid buffer) were run between each sample to reduce carry-over. The column was kept at a constant temperature of 50 °C.

The data was acquired using an easy spray source operated in positive mode with spray voltage at 2.445 kV, and the ion transfer tube temperature at 250 °C. The MS was operated in DIA mode. A scan cycle comprised a full MS scan (m/z range from 350 to 1650), with RF lens at 40%, AGC target set to custom, normalised AGC target at 300%, maximum injection time mode set to custom, maximum injection time at 20 ms, microscan set to 1 and source fragmentation disabled. MS survey scan was followed by MS/MS DIA scan events using the following parameters: multiplex ions set to false, collision energy mode set to stepped, collision energy type set to normalized, HCD collision energies set to 25.5, 27 and 30%, orbitrap resolution 30,000, first mass 200, RF lens 40%, AGC target set to custom, normalized AGC target 3000%, microscan set to 1 and maximum injection time 55 ms. Data for both MS scan and MS/MS DIA scan events were acquired in profile mode.

### Mini-bulk and single cell proteomics sample preparation

Single neutrophils and mini-bulk samples (500 cells) were sorted into fresh 384 well plates (Thermo Scientific™ Armadillo PCR Plate, 384-well, #12657516) each well containing 1 μL of master mix. For 0 cell samples, no cells were sorted into the respective master mix-containing wells. The master mix contained 0.2% n-dodecyl-ß-D-maltoside (DDM, #D4641-500MG, Sigma Aldrich, Germany), 100mM triethylammonium bicarbonate (TEAB, #17902-500ML, Fluka Analytical, Switzerland), and 3 ng/μL trypsin (Trypsin Gold, #V5280, Promega, USA) in ultra-pure water. Directly after sorting, samples were stored at -80°C. Just prior to lysis and digestion, plates were thawed and centrifuged for 3min at 300xg at 4°C. Lysis and digestion was performed similarly to earlier works^16^. In brief, plates were placed in the CellenONE X1 (Cellenion, Lyon, France, #F00C) at 50°C and 85% humidity to limit evaporation. This is followed by incubation for 2 h at 50°C at 85% relative humidity inside the instrument. Samples were kept hydrated every 15 min by automated addition of 500 nL ultrapure to each well. After 30 min of incubation, an additional 500nL of 3 ng/μL trypsin were added which replaces one hydration step. After lysis and digestion, 2.5 μL of 0.1% trifluoro acetic acid (TFA, Thermo Fisher Scientific, #28903) with 5% dimethyl sulfoxide (Avantor, #83673.230) were added to the respective wells for quenching and storage. Plates were then stored at -20°C before LC-MS/MS measurement.

### Mini-bulk and single cell proteomics mass spectrometry

All samples were analyzed using a Vanquish Neo UHPLC (Thermo Fisher Scientific, #VN-S10-A-01) operated in trap-and-elute mode and coupled to an Orbitrap Astral mass spectrometer (Thermo Fisher Scientific, #BRE725600) equipped with a FAIMS Pro Duo interface (ThermoFisher Scientific, #OPTON-20068). Analyte separation was performed at 50°C using a 25 cm x 75 μm C18 UHPLC packed emitter column (Ion Opticks Pty Ltd, # AUR3-25075C18-TS, Collingwood, Australia) connected to the mass spectrometer via an EASY-Spray^TM^ ion source (Thermo Fisher Scientific, ES081). Before separation, peptides were trapped on a 5mm PepMap^TM^ Neo trap cartridge (Thermo Fisher Scientific, #174500) using a flow of 10 µL/min of 0.1% TFA. Loading was done using combined control of flow (10µL/min) and pressure (max. 800 bar). Loading volume was set to automatic. Samples were cooled to 7°C in the autosampler and covered with a silicone mat. 5 µL were injected per sample to ensure complete aspiration of the entire sample volume using 0.2µL/s draw speed and 2s draw delay with bottom detection on. For separation, 0.1% formic acid in LC-MS/MS grade water (Thermo Fisher Scientific, # 10188164) as solvent A and 0.08% formic acid in 80% acetonitrile (Thermo Fisher Scientific, #10118464) as solvent B were used. After sample loading, peptides were eluted using the following gradient as previously described^20^ to achieve a throughput of roughly 50 samples per day: 0.0min 450nL/min 1%B - 0.1min 450nL/min 4%B – 1.9min 450nL/min 12%B – 2.0min 200nL/min 12%B – 12.0min 200nL/min 22.5%B – 19.5min 200nL/min 40%B – WASH PHASE – 22.0min 300nL/min 99%B – 25.0min 300nL/min 99%B.

An electrospray voltage of 1.9 kV was applied for peptide ionization with a static carrier gas flow of 3.5 L/min. The ion transfer tube temperature was set to 280°C, expected peak width to 6s, the default charge state to 2 and advanced peak determination was enabled. Total method length was set to 27min. MS1 were recorded in positive polarity in the orbitrap at 240,000 resolution for a scan range of 400-800 m/z. The AGC target was set to 500% (= 5E6 absolute AGC target) with 100ms as maximum injection time. A single FAIMS CV of -48 V was used and profile chosen as data type. RF lens (%) was kept at 45.

Fragmentation spectra were recorded using DIA with 20 m/z isolation windows without any overlap and window placement optimization on. DIA MS2 spectra were recorded in the Astral mass analyzer for a precursor range of 400-800 m/z and a scan range of 150-2000 m/z. Collision energy was set to 25% NCE and the AGC target to 800% (=80,000 absolute AGC target). The maximum injection time was set to 40ms for mini-bulk and 80ms for single and 0 cell samples. Centroid was chosen as data type and 0.6s selected as loop control. DIA window type was set to Auto and mode to m/z range.

### Spectronaut analysis

The bulk and mini-bulk were searched with Spectronaut^49^ 19.7, the single cell was searched with Spectronaut 19.4. For all 3 searches the default parameters were altered to increase stringency similar to Baker et al^50^. The details are specified in Table 1. All searches were performed using directDIA against a human SwissProt + isoforms database (November 2024) and an immune cell specific contaminant fasta file. The fasta files are included in the PRIDE submissions. No variable modifications were included in the searches.

### Spectronaut parameters

**Table 1.** Spectronaut parameters

| Parameter | Value |
| --- | --- |
| Precursor Qvalue Cutoff | 0.01 |
| Protein Qvalue Cutoff<br>(experiment) | 0.01 |
| Protein Qvalue Cutoff (run) | 0.01 |
| Precursor PEP Cutoff | 0.15 |
| Protein PEP Cutoff | 0.15 |
| Protein LFQ Method | Quant2.0 |
| Quantity MS method (bulk) | MS2 |
| Quantity MS method (mini-bulk<br>& single cell) | MS1 |
| Cross-Run Normalisation | Off |
| Major (Protein) Grouping | Protein Group ID |
| Major Group Quantity | Peptide sum |
| Major Group Top N | Off |
| Minor (Peptide) Grouping | By Stripped Sequence |
| Minor Group Quantity | Median precursor quantity |
| Minor Group Top N | Off |

### Copy number calculation

For all data in figure 1, estimated protein copy numbers were calculated using the healthy control participants. Copy numbers were calculated from the mass spectrometry-based intensity data using the proteomic ruler^27^. The median copy numbers for bulk, mini-bulk and SCP were calculated using the bulk estimations but for proteins identified with each methodology.

### Proteomic data normalisation

For mini-bulk the intensity based absolute quantification (iBAQ) was used and divided by the median intensity across each sample to generate the relative iBAQ (riBAQ) for all proteins across all samples. For SCP, due to issues in the iBAQ calculation with Spectronaut 19.4, an iBAQ like measure was calculated. In brief, the protein intensity was divided by the protein molecular weight and multiplied by the median molecular weight of all detected proteins (47,485Da). Across each sample this molecular weight corrected intensity was divided by the sample median producing a riBAQ like quantitation.

### Coefficient of variation (cv) calculations

The CVs were calculated using the proteomicsCV^51^ package v0.4.0. The raw intensities were fed to the package after performing the previously stated riBAQ normalization.

### Bulk and mini-bulk differential expression analysis

The differential expression analysis was perform in R using the Bioconductor package limma^52^ v3.60.6 using lmFit with the method=’robust’ parameter and eBayes with the ‘robust=TRUE’ parameters. Q values were calculated using the Bioconductor package qvalue v2.36.

### Mini-bulk volcano plot filters

For all mini-bulk volcano plots only proteins that were detected in at least 2 independent replicates in both conditions being compared were considered. Any proteins detected in 1 replicate or less in any conditions were filtered out.

### Significant proteins

For the mini-bulk analysis, proteins were considered to be significantly changed in abundance if their q-value<0.05. For single cell proteomics they were considered significant if the Bonferroni adjusted p-values were <0.05.

### Mini-bulk gene ontology (go) analysis

The GO analysis was performed using WebGestalt^53^ with a minimum number of analytes per category set to 8, Significance Level set to FDR 0.05, and weighted set cover, affinity propagation and k-Mediods enabled. The background was set to all proteins detected in at least two replicates in both NDN CD10+ and LDN CD10-, and two replicates in the TAN and LDN CD10-. Proteins that were significantly increased in abundance in LDN CD10-were analysed against the Biological Process and Cellular Component non-redundant databases.

### Principal component analysis

All mini-bulk based PCAs were analysed in R using only proteins that were identified in all samples and after log10 transformation using the prcomp function part of the stats v4.4.1 package.

### Mini-bulk sample filtering

One control sample had 10-fold lower number of proteins and peptides identified and was a clear outlier. This sample was filtered from the posterior analysis.

### Single cell proteomics cell filtering

The 0 cell runs were used as QC for filtering. A threshold of >1.75-fold higher than the maximum number of identifications in the 0 cell runs (Extended Data Fig. 5A) was set and rounded up to 400 proteins. A total of 277 cells were analysed after filtering.

### Seurat pipeline

The single cell proteomics data was analysed using Seurat^54^ v.5.2.1. The intensity data were normalised as described above and this normalisation was used in Seurat. The min.features were set to 400 (filtering cells with less than 400 proteins identified) and FindVariableFeatures used nfeatures = 2000. Total number of principal components (npcs) was set to 35. The data were batch corrected using the R package harmony v.1.2.3 and grouping the data by patient. FindNeighbours() used the harmony input with a total of 35 dimensions, FindClusters() used a resolution of 1.45. The number of clusters were determined using the ElbowPlot() method and set to 7.

### Single cell population markers

For the population markers the FindMarkers() function in Seurat was used. A fold-change threshold of 0.25, logistic regression as the selected method, and a Bonferroni corrected p-value < 0.05 was considered significant.

### Single cell protein family significance testing

All protein family significance was calculated using the sum of the intensity of all proteins within the family and calculated using Welchs T test in R. A p-value <0.05 was considered significant

### Single cell proteomics data visualisations

The doplots and UMAP were all generated with Seurat. The sankey diagram was generated with Sankeymatic. The violin plots were generated in R using ggplot2 v3.5.1

## ACKNOWLEDGMENTS

Flow cytometry data were generated with support from the QMRI and IRR Flow Cytometry and Cell Sorting Facility, within the University of Edinburgh. This work was funded by a CRUK Cancer Immunology project award C62207 (SRW), Wellcome Trust Senior Clinical Fellowship awards 098516, 209220 (SRW), Discovery science award 225778 (SRW) and a Wellcome Trust Clinical Research Career Development Fellowship award 224637/Z/21/Z22 (EW). This work was also supported by the infrastructure funding fourth call 2022/01 of the Austrian Research Promotion Agency (KM) and the project LS20-079 of the Vienna Science and Technology Fund (KM), the P35045-B project (grant DOI 10.55776/P35045) and the project PAT4142423 (Grant DOI 10.55776) of the Austrian Science Fund (KM). We thank the Protein Chemistry Facility and acknowledge the VBCF for instrument access. We also thank the Lothian NRS bioresource, the consultant neuro-anesthetists and registrars of the department of clinical neurosciences, NHS Lothian for assistance in recruiting, consenting and obtaining samples from patients with Glioblastoma. For the purpose of open access, the author has applied a CC BY public copyright license to any Author Accepted manuscript version from this submission.

## AUTHOR CONTRIBUTIONS

Conceptualization: SRW; Methodology: AJB, PS, LR, RLM, LC, SMP, KM; Investigation: AJB, PS, LR, PC, GVS, AZ, MAS-G, ERW, IL, AJMH, IA, AB, AM, SR GMM, HM, CB, RG, FAM, XRI, NP, MR, PMB; Funding acquisition: SRW, KM; Supervision: SRW, PS; Writing – original draft: SRW, AJB, PS; Writing – review & editing: SRW, PS, AJB, GVS, DAC, RLM, KM

## CORRESPONDING AUTHORS

Correspondence to Sarah R. Walmsley, Karl Mechtler or Alejandro J. Brenes.

## DECLARATION OF INTERESTS

SMP is a founder, consultant and shareholder of Trogenix Ltd, a biotech that is working on advanced therapies for cancers of unmet need. He is an inventor on patents owned by the University of Edinburgh that have been licensed to Trogenix.

## Supplementary Information

**Extended Data Fig. 1.**
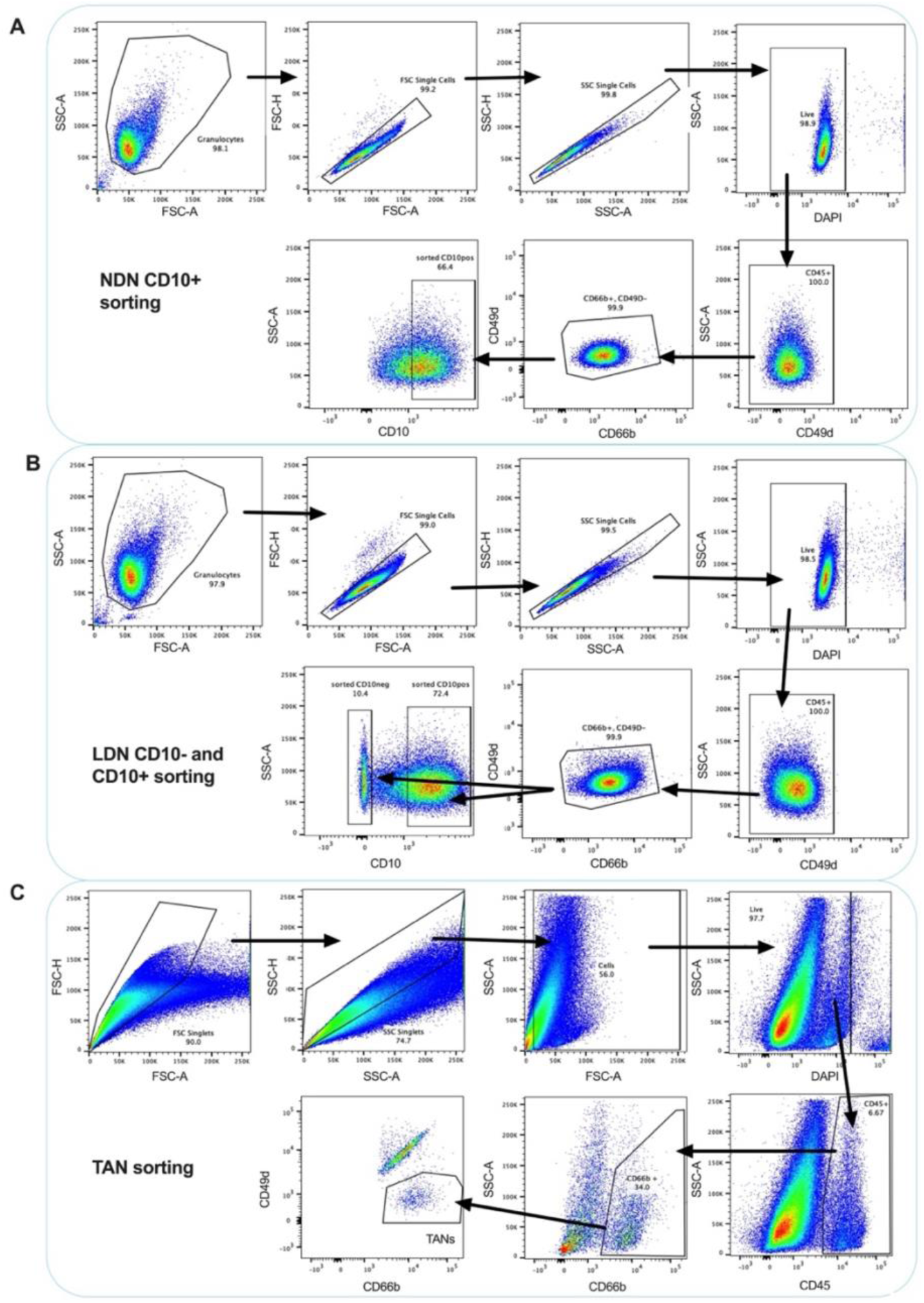
Gating strategy of neutrophils sorted from GBM blood and tumours. Representative dot plots illustrating the gating strategy of neutrophils sorted from circulating GBM NDN (A), LDN (B) and tumours post enzymatic digestion (C). 500 cells GBM NDN CD10+, LDN CD10-, LDN CD10+ and TAN subpopulations in addition to healthy control NDN (n=6 for each subpopulation) were sorted into 384 well plates for the mini-bulk proteomic. For SCP analyses, single TANs were sorted into 384 well plates (n=330). NDN were gated on live CD45+, CD66b+, CD49-, CD10+ and LDN neutrophils were gated on live CD45+, CD66b+, CD49-,CD10- (immature) and live CD45+, CD66b+, CD49d-,CD10+ (mature) cells whereas TANs were gated on live CD45+, CD66b+, CD49d-for both mini-bulk and SCP analysis.

**Extended Data Fig. 2.**
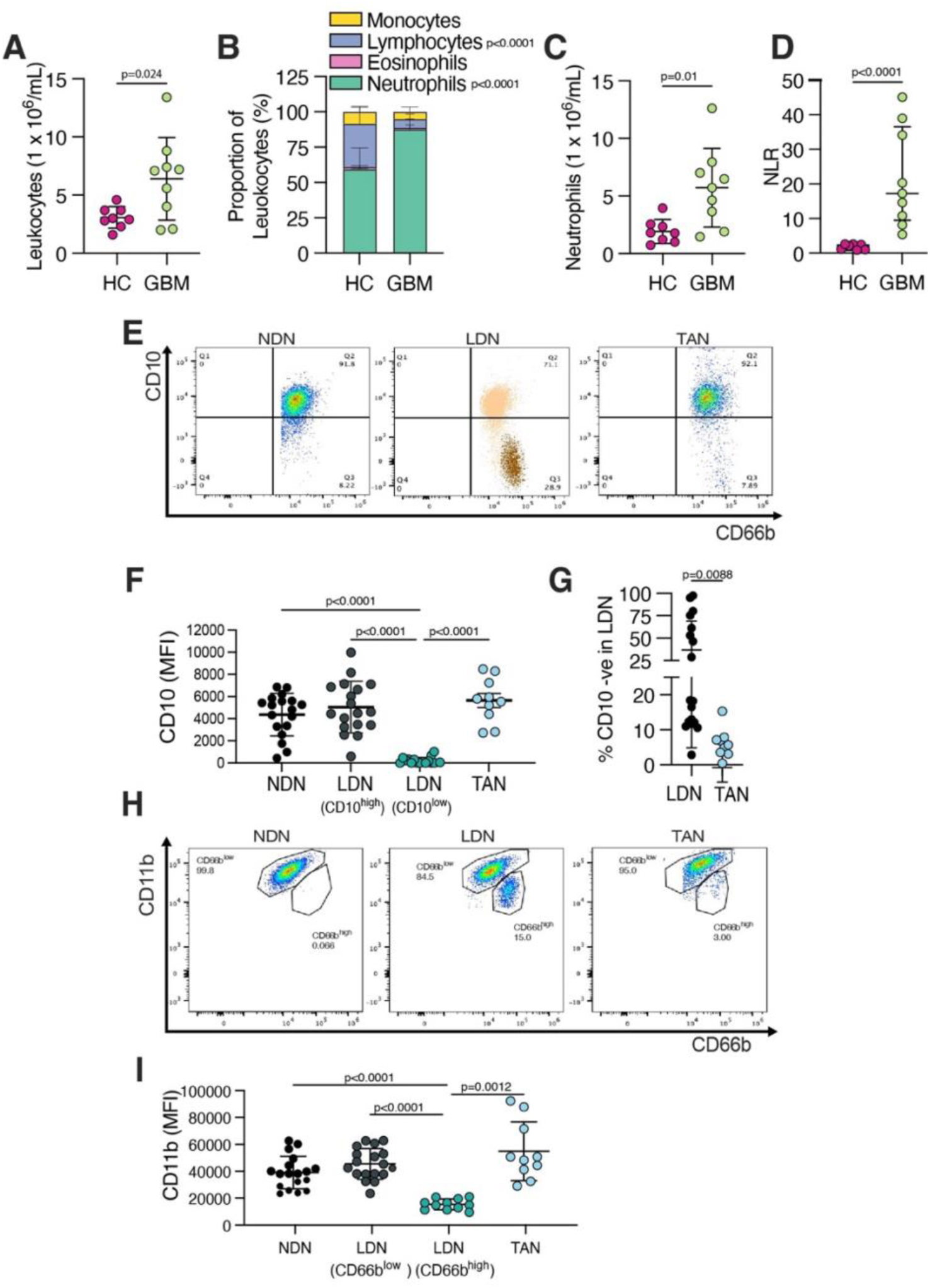
Neutrophil expansion and presence of an immature circulating neutrophil population. (A-D) Flow cytometry analysis was performed on whole blood from healthy donors (HC, n = 8) and GBM patients (n = 9) and total leukocyte count **(A)**, leukocyte proportions **(B)**, total neutrophil counts **(C)** and neutrophil to lymphocyte ratio (NLR) **(D)** quantified. **(E)** Representative CD10 vs CD66b dot plots of circulating normal density neutrophils (NDN) (left), low density neutrophils (LDN) (middle) and tumour associated neutrophils (TAN) in GBM (right). (**F**) Surface expression of CD10 expressed as median fluorescence intensity (MFI), NDN n=18, LDN CD10^high^ n=18, LDN CD10^low^ n=17, TAN n=10. (**G**) Percentage of cells with CD10-expression in LDN (n=18) and TAN (n=9). (**H**) Representative CD11b vs CD66b plots of circulating NDN, LDN and TAN in GBM gated on live, CD66b+, CD49d-expression. **(I)** Surface expression of CD11b expressed as MFI, NDN n=18, LDN CD10^high^ n=18, LDN CD10^low^ n=11, TAN n=10. Data represent mean ± SD. P-values were calculated with Welch’s t-test (A, C, D, G), two-way ANOVA with Sidak’s multiple comparison test on transformed data to Arcsin (B), Brown-Forsythe and Welch ANOVA tests (F, I).

**Extended Data Fig. 3.**
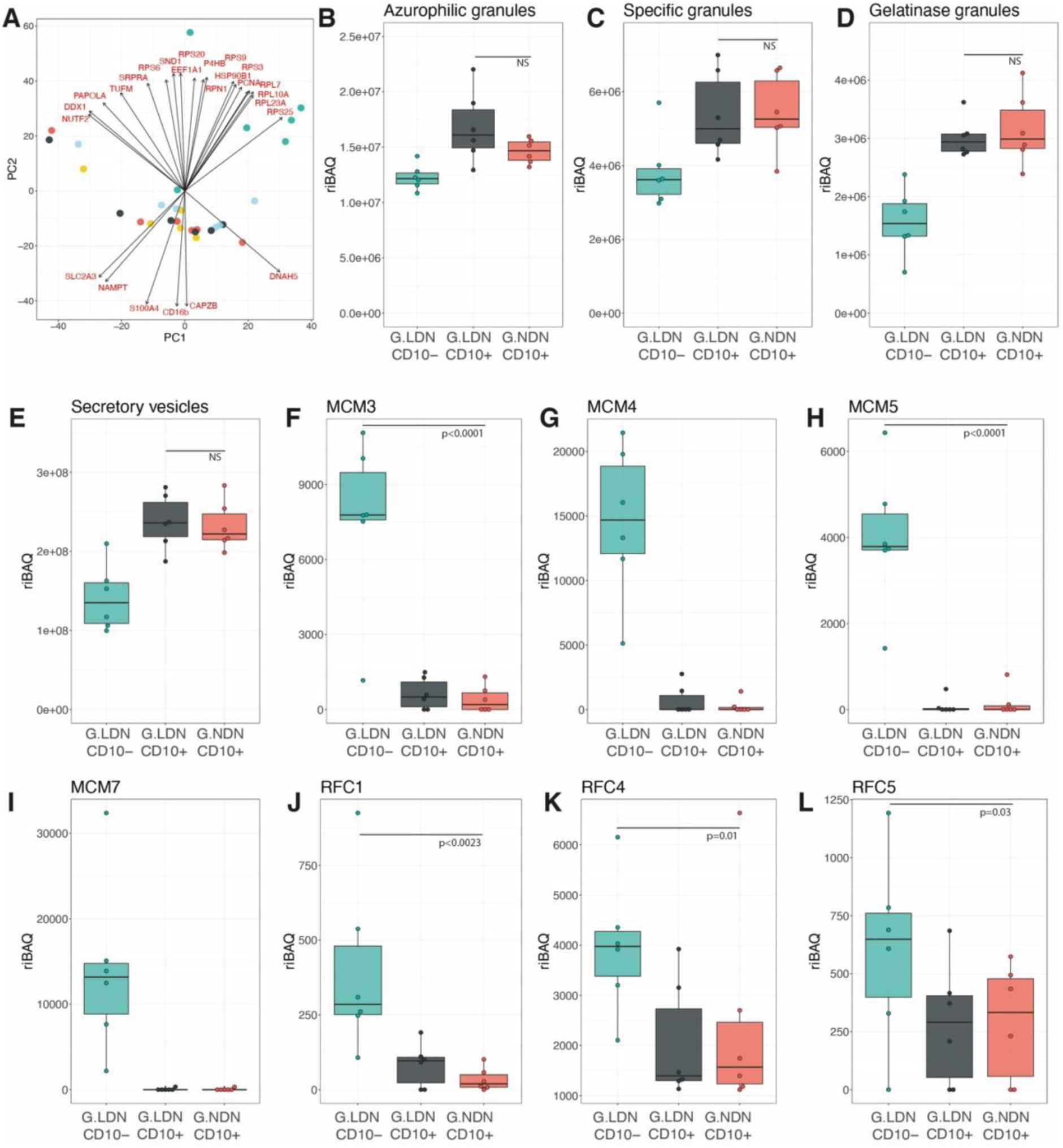
Granule and DNA replication proteins in mature (CD10+) and immature (CD10-) neutrophil proteomes. (A) PCA plot showing the top 25 protein components for all mini-bulk samples. Boxplots (n=6 across all conditions) showing the riBAQ for (B) all azurophilic granules (C) all specific granules, (D) all gelatinase granules, (E) all secretory vesicles, (F) Minichromosome Maintenance Complex Component 3 (MCM3), (G) Minichromosome Maintenance Complex Component 4 (MCM4), (H) Minichromosome Maintenance Complex Component 5 (MCM5), (I) Minichromosome Maintenance Complex Component 7 (MCM7), (J) Replication Factor C Subunit 1 (RFC1), (K) Replication Factor C Subunit 4 (RFC4) and (L) Replication Factor C Subunit 5 (RFC5) across LDN CD10-, LDN CD10+, NDN CD10+. For all boxplots the top and bottom hinges represent the 1st and 3rd quartiles. The top whisker extends from the hinge to the largest value no further than 1.5 × interquartile range (IQR) from the hinge; the bottom whisker extends from the hinge to the smallest value at most 1.5 × IQR of the hinge. All p-values were calculated with limma using Empirical Bayes statistics for differential expression.

**Extended Data Fig. 4.**
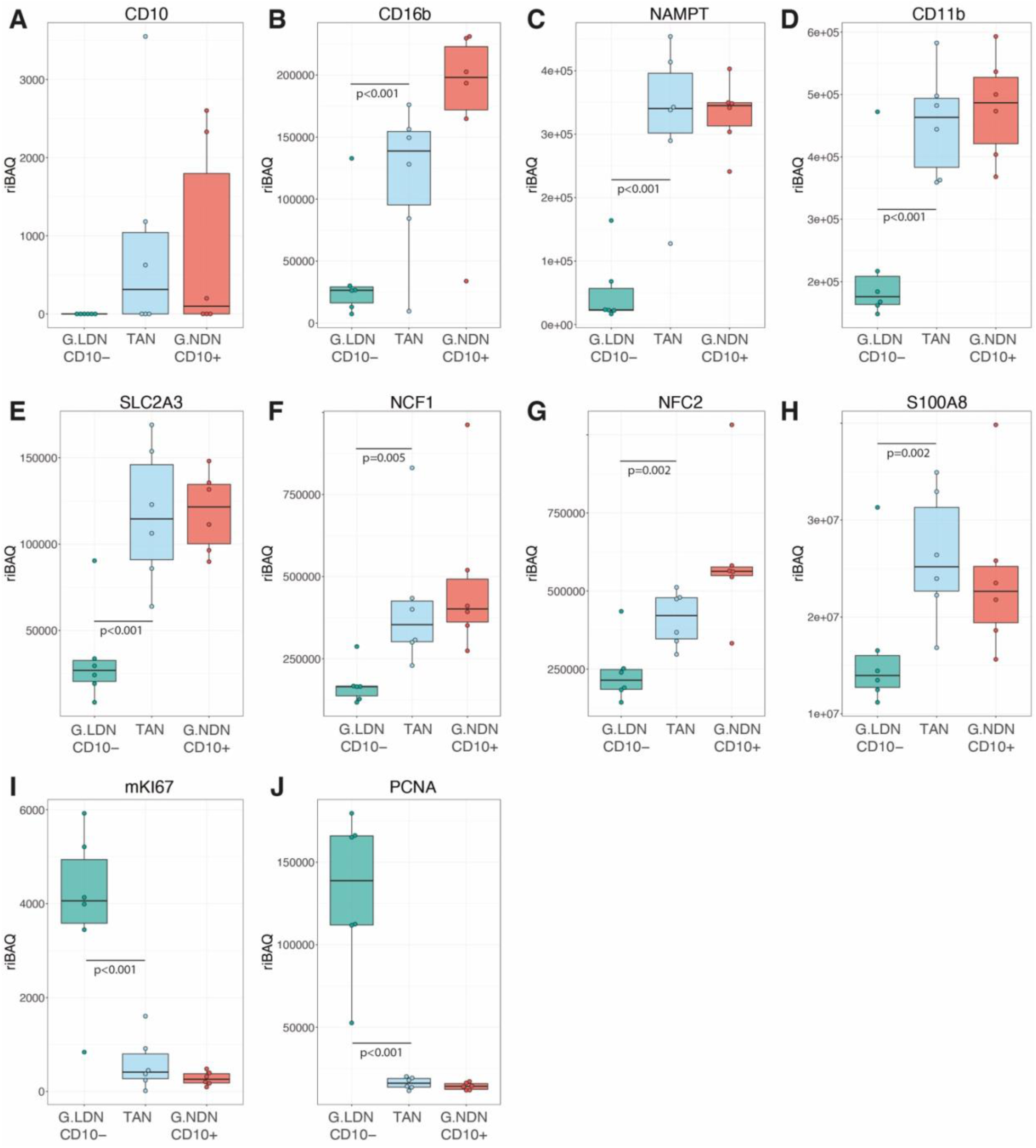
Maturity and proliferation markers across NDN CD10+, LDN CD10- and TANs. Boxplots (n=6 across all conditions) showing the riBAQ for (A) CD10 (B) CD16b, (C) Nicotinamide Phosphoribosyltransferase (NAMPT), (D) CD11b, (E) Solute Carrier Family 2 Member 3, GLUT 3 (SC2A3), (F) Neutrophil Cytosolic Factor 1 (NCF1), (G) Neutrophil Cytosolic Factor 2 (NCF2), (H) S100 Calcium Binding Protein A8 (S100A8), (I) Marker Of Proliferation Ki-67 (mKI67) and (J) Proliferating Cell Nuclear Antigen (PCNA) across LDN CD10-, NDN CD10+ and TANs. For all boxplots, the top and bottom hinges represent the 1st and 3rd quartiles. The top whisker extends from the hinge to the largest value no further than 1.5 × interquartile range (IQR) from the hinge; the bottom whisker extends from the hinge to the smallest value at most 1.5 × IQR of the hinge. All p-values were calculated with limma using Empirical Bayes statistics for differential expression.

**Extended Data Fig. 5.**
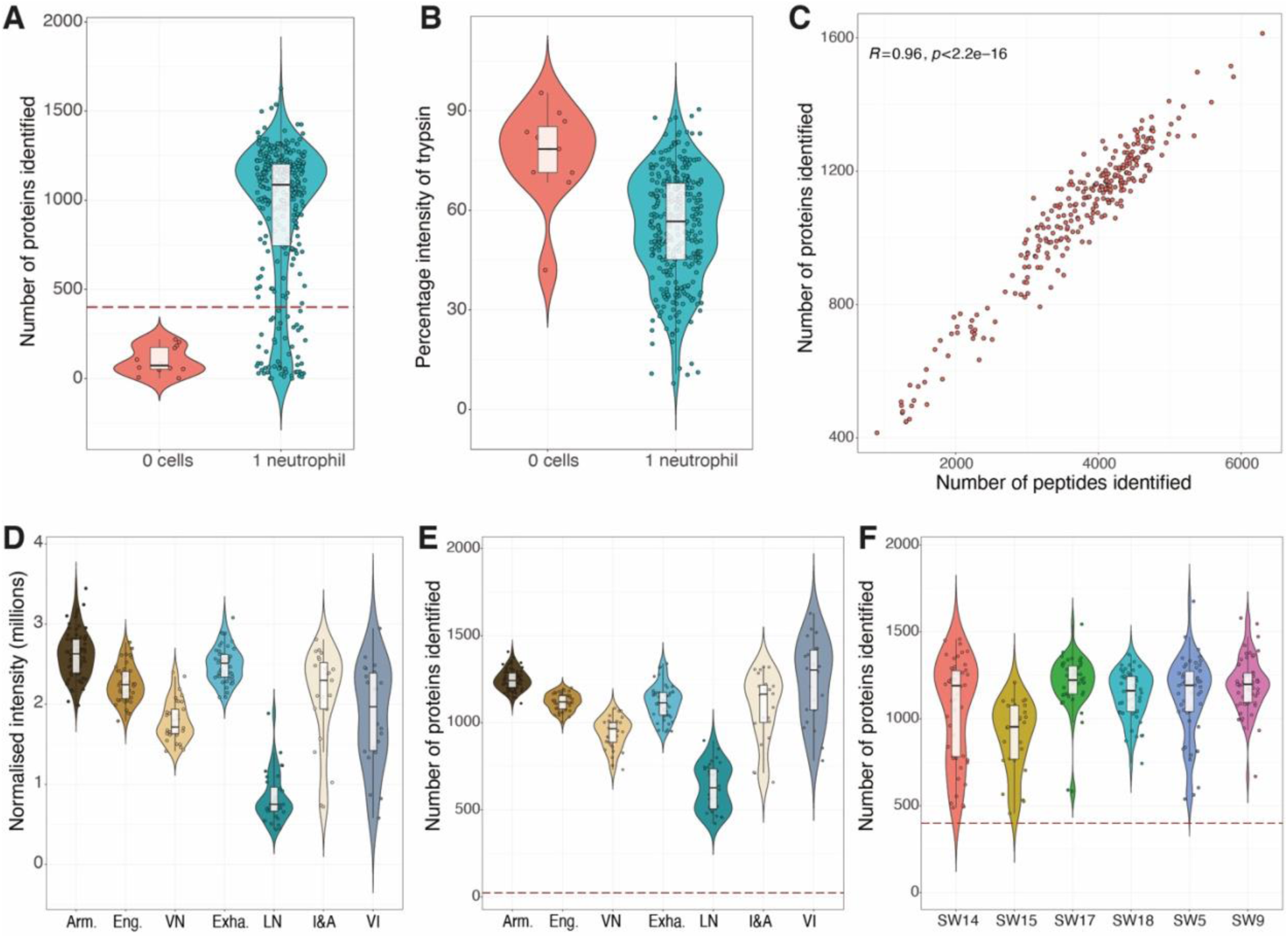
Single cell proteomics quality control. (A) Boxplots showing the number of proteins identified in the 0 cell (n=12) and the single neutrophil runs (n=330). The red dotted line represents 400 proteins. (B) Boxplots showing the percentage of the total intensity represented by trypsin in the 0 cell (n=12) and the single neutrophil runs (n=330). (C) Scatter plot showing the number of proteins and peptides identified in each individual cell (n=277). Boxplots showing the (D) total normalized intensity and the (E) number of proteins identified across all 7 neutrophil clusters. Armed (n=76), Engaged (n=46), Vital NETs (n=33), Exhausted (n=47), Lytic NETs (n=32), Immunosuppressive and angiogenic (n=23) and Vascular Immature (n=22). (F) Boxplot showing the number of proteins identified in each single neutrophil organized by patient. SW14(n=48), SW15(n=28), SW17(n=45), SW18(n=51), SW5(n=50), SW9(n=55). For all boxplots, the top and bottom hinges represent the 1st and 3rd quartiles. The top whisker extends from the hinge to the largest value no further than 1.5 × interquartile range (IQR) from the hinge; the bottom whisker extends from the hinge to the smallest value at most 1.5 × IQR of the hinge.

**Extended Data Fig. 6.**
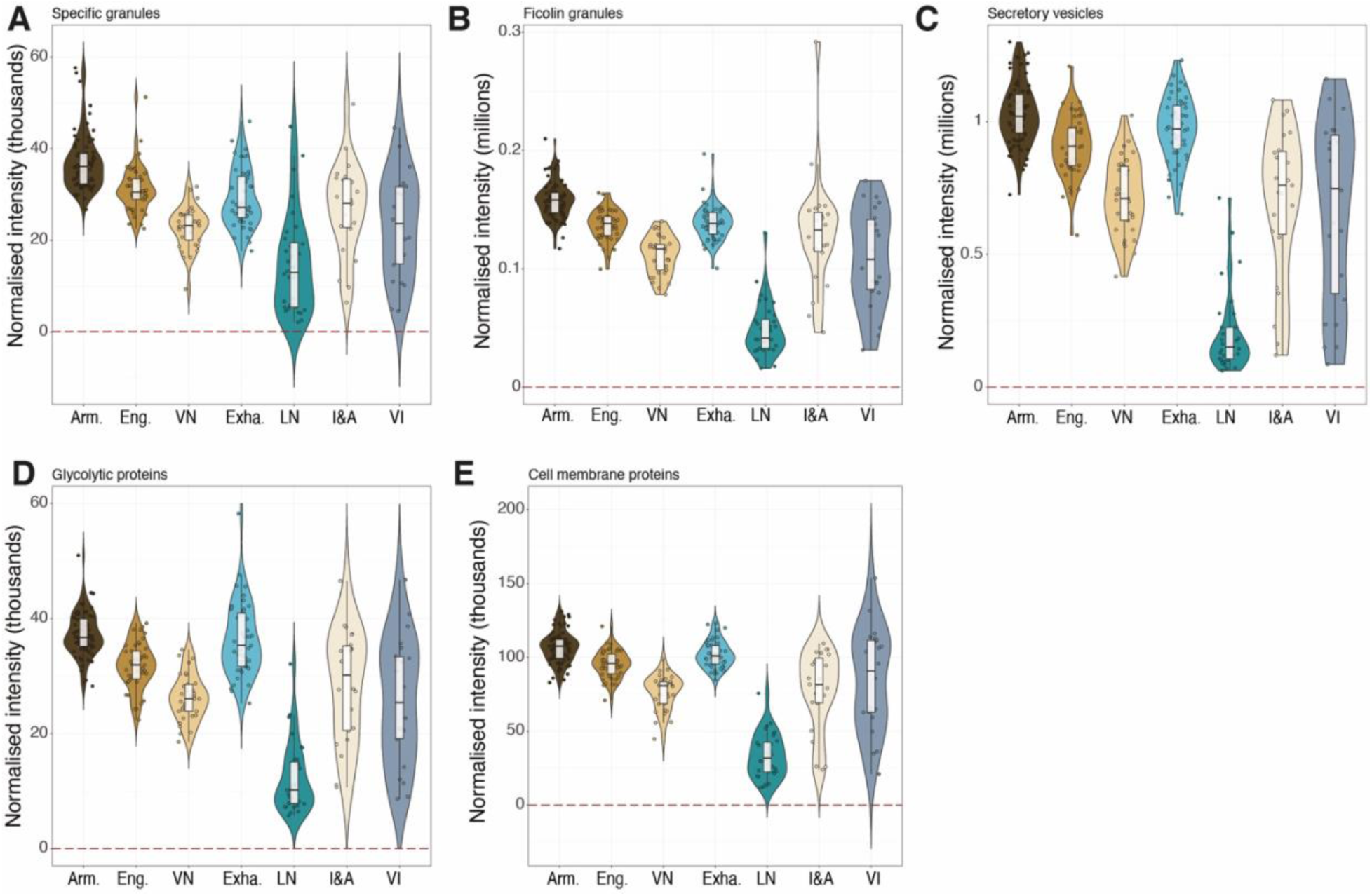
Lytic NETs reductions in granules, metabolism and cell surface proteins. Boxplots showing the sum of the normalized intensity for all proteins labelled as (A) specific granules, (B) ficolin granules, (C) secretory vesicles, (D) glycolytic proteins and (E) cell membrane proteins across all 7 neutrophil clusters. Across all boxplots Armed (n=76), Engaged (n=46), Vital NETs (n=33), Exhausted (n=47), Lytic NETs (n=32), Immunosuppressive and angiogenic (n=23) and Vascular Immature (n=22). For all boxplots the top and bottom hinges represent the 1st and 3rd quartiles. The top whisker extends from the hinge to the largest value no further than 1.5 × interquartile range (IQR) from the hinge; the bottom whisker extends from the hinge to the smallest value at most 1.5 × IQR of the hinge.

